# Tempered IL-2 Signals Program PD-1 Checkpoint Blockade-Responsive Stem-like Exhausted T Cells During Priming

**DOI:** 10.1101/2023.05.25.541936

**Authors:** Vandana Kalia, Jim Reed, Asheema Khanna, Ryma Toumi, Hanxi Xiao, Thomas Pulliam, Rucha Deo, Paul Nghiem, Surojit Sarkar

## Abstract

Stem-like progenitor exhausted CD8 T cells are critical for maintaining long-term resistance during chronic infections and cancer, and represent an important checkpoint blockade immunotherapy target for functional reinvigoration and disease control. Hence, there is vigorous interest in understanding the ontogenesis of TCF-1^Hi^ stem-like exhausted CD8 T cells, and in defining the signals that promote their development. Here, we show that virus-specific TCF1^Hi^GzmB^Lo^ CD8 T cells develop during early stages of chronic viral infection, and are progressively deprogrammed from the stem-like lineage towards terminal differentiation by strong IL-2 signals. In vivo fate-tracking studies show that strong T cell-intrinsic IL-2 signals through the high affinity heterotrimeric IL-2Rα/β/γ receptor drive skewed development of terminally differentiated TCF-1^Lo^GzmB^Hi^ cells, which largely die in chronic antigenic environment. In contrast, tempered IL-2 signals through the intermediate affinity IL-2Rβ/γ heterodimer, or delayed priming in diminished IL-2 milieu support preferential development of TCF-1^Hi^ stem-like exhausted CD8 T cells, capable of long-term persistence and potent responsiveness to PD-1 therapy in later stages of chronic viral infection. In human tumors as well, single cell RNA-seq analyses of tumor infiltrating lymphocytes from melanoma, human papillomavirus+ head and neck cancer and lung cancer patients revealed an inverse relationship between IL-2 signaling signature and T cell stemness. Moreover, melanoma patients with enriched IL-2 signaling signature showed poor responses to checkpoint blockade immunotherapy. Collectively, these findings bear relevance to clinical immunotherapies of chronic viral infections and cancers, and support exogenous IL-2 signal manipulation prior to therapy as a strategy for augmenting long-term clinical outcomes.

**ONE SENTENCE SUMMARY:** Programming of exhausted T cell fates by differential IL-2

## INTRODUCTION

Exhausted virus-specific CD8 T cells critically restrain pathogen outgrowth despite functional impairment ^1^. This property is ascribed to a small subset of stem-like CD8 T cells, which have been recently identified within the exhausted CD8 T cell pool of chronic viral infections as well as cancer ^2–12^. Distinguishable from terminally exhausted CD8 T cells by higher expression of T cell factor-1 (TCF-1) and lower expression of inhibitory receptors, the stem-like exhausted CD8 T cells are capable of self-renewal, and have been shown to continually seed the transient pool of functional effector cells for mediating host-virus stand-off ^8, 10, 13, 14^. Importantly, TCF-1^Hi^ stem-like exhausted CD8 T cells undergo vigorous expansion and functional reinvigoration in response to PD-1 checkpoint blockade immunotherapy (CBI), and are crucial for reducing viral loads and tumor burdens post-therapy ^3–5, 7, 9, 15^. TCF-1^Hi^ stem-like CD8 T cells have also been delineated in a multitude of human cancers such as head and neck cancer ^16^, melanoma ^7, 17^, lung cancers ^18–21^ and hepatocellular carcinoma (HCC) ^22^. Hence, inducing stem-like T cells is a highly desirable goal in chronic infections and cancers for mediating long-term disease control, and for the therapeutic success of PD-1 therapy as well as adoptive T cell transfer (ACT) therapeutic interventions. While our understanding of the transcriptional regulation of T cell exhaustion is quite extensive with several known factors such as TOX, Blimp-1, E2A, T-bet, Eomes, BACH-2, BATF, IRF-4 and Myb ^4, 8, 14, 23–34^, we know very little about the origins and signals that drive the initial generation and subsequent development of TCF-1^Hi^ stem-like CD8 T cells in the context of chronic antigenic settings.

## RESULTS

### TCF-1^Hi^ stem-like CD8 T cells are distinguishable during early stages of priming in chronic viral infection, and correlate with lower expression of IL-2R heterotrimer

Key features of T cell exhaustion manifest early during CD8 T cell responses to chronic LCMV infection (days 5-8 after infection)^8, 11–13, 35–37^, and it is proposed that TCF-1^Hi^ GzmB^Lo^ stem-like and TCF-1^Lo^ GzmB^Hi^ terminally differentiated CD8 T cell lineages are programmed during initial priming and expansion. We used carboxyfluorescein succinimydyl ester (CFSE) labeled D^b^GP33-specific CD8 T cells to analyze the expression patterns of TCF-1 and GzmB *vis a vis* cell division early after activation (days 2.5-3.5). As shown in Fig 1A, TCF-1 expression progressively decreased, whereas GzmB expression conversely increased with increasing rounds of cell division, thus leading to a clear heterogeneity within the early virus-specific CD8 T cell pool. Both TCF-1^Hi^ GzmB^Lo^ and TCF-1^Lo^ GzmB^Hi^ CD8 T cell subsets were evident as early as days 3.5-5.0 after LCMV_Cl-13_ infection in both TCR-transgenic D^b^GP33-specific P14 CD8 T cells at low or high precursor frequencies (Extended Fig 1A) as well as in endogenous tetramer+ D^b^GP33, D^b^GP276 and D^b^NP396-specific CD8 T cells (Fig 1B). Consistent with previous reports of increased terminal exhaustion and clonal deletion of NP396-specific CD8 T cells, we noted lower proportions of stem-like cells in NP396-specific CD8 T cells during early stages of infection (day 5.5), followed by clonal deletion by about day 8 after infection. As in the case of stem-like exhausted CD8 T cells isolated from late stages of chronic infection, the TCF-1^Hi^ GzmB^Lo^ stem-like cells isolated during early stages of infection showed preferential enrichment in secondary lymphoid organs (spleen and lymph nodes) compared to peripheral lung and liver sites (Extended Data Fig 1B).

**FIGURE 1.**
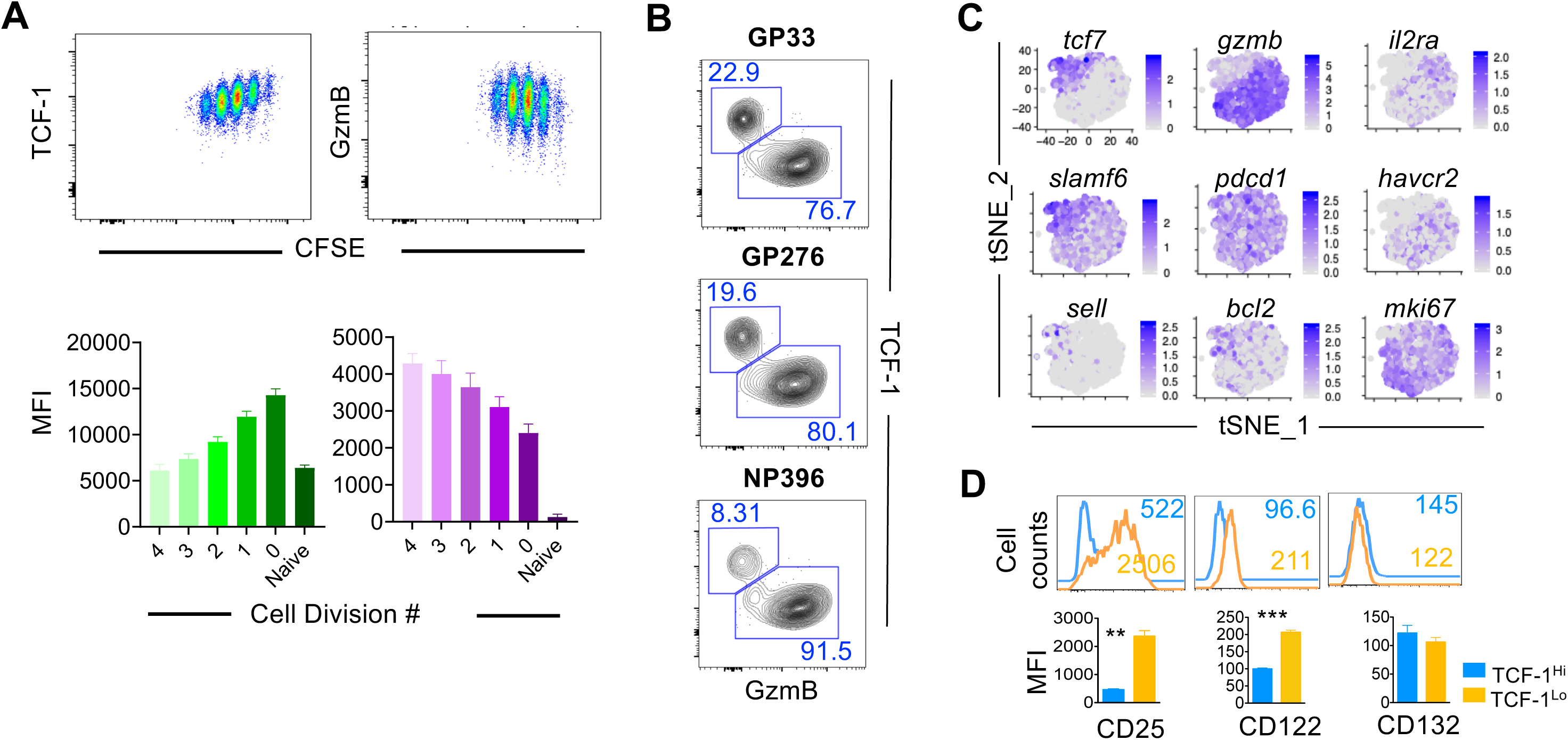
TCF-1^Hi^ stem-like CD8 T cell lineage is preferentially associated with reduced IL-2R expression during T cell priming in chronic viral infection. **(A)** D^b^GP33-specific CFSE labeled P14 CD8 T cells were stimulated with GP33 peptide loaded C57Bl/6 splenocytes and analyzed for CFSE dilution and indicated markers at day 2.5 after activation. **(B)** Naïve C57Bl/6 mice were infected with LCMV_Cl-13_ and splenocytes were analyzed for expression of TCF-1 and GzmB in GP33, GP276 and NP396 tetramer+ CD8 T cells. **(C)** scRNA-seq data analysis of markers associated with stem-like CD8 T cells and IL-2Rα in chronic LCMV infection. Single-cell transcript levels of *tcf7*, *gzmb*, *il2ra*, *slamf6*, *pdcd1*, *havcr2*, *sell*, *bcl2* and *mki67* are illustrated in t-SNE plots from LCMV-specific CD8 T cells 4.5 days after LCMV_Cl-13_ infection (Transcript levels are color-coded: grey, not expressed; purple, expressed). **(D)** WT D^b^GP33-specific P14 cells (2.5×10^3^, low dose, endogenous precursor frequencies) were adoptively transferred into naïve B6 mice, which were subsequently infected with LCMV_Cl-13_. Flow-cytometry plot of TCF-1 and GzmB co-expression in splenic P14 CD8 T cells are shown at day 5.5 post-infection. Histograms and bar graphs depict levels of CD25, CD122 and CD132 expression on TCF-1^Hi^ (Light blue) or TCF-1^Lo^ (Gold) gated P14 T cells. Numbers represent MFI of expression of respective markers. Data are representative of 4 independent experiments (mean ± SEM) with at least 3 mice per group.

Towards gaining insight into the exhausted T cell fate programming signals, we conducted scRNA-seq data analysis of virus-specific CD8 T cells isolated from early stages of murine infection with chronic LCMV_Cl-1336_. Consistent with the notion of early programming of the stem-like CD8 T cell lineage, T-distributed stochastic neighbor embedding (t-SNE) analysis discerned TCF-1^Hi^ cells, which also expressed higher levels of Slamf6, Bcl-2 and CD62L, and inversely with proliferation marker Ki-67, inhibitory receptor Tim3 and effector molecule GzmB (Fig 1C)^8, 36^. Intriguingly, we noted an inverse expression pattern of TCF-1 with IL-2Rα or CD25 at both mRNA and protein levels (Fig 1C, 1D, Extended Data Fig 1A). Compared to TCF-1^Lo^ GzmB^Hi^ cells, TCF-1^Hi^ GzmB^Lo^ cells expressed lower levels of IL-2Rα (CD25) − a component of the IL-2Rα/β/γ heterotrimer that imparts high affinity IL-2 binding – as well as the common IL-2Rβ-chain (CD122), with largely similar expression of the common IL-2Rγ-chain (CD132) (Fig 1D). The TCF-1^Hi^ CD25^Lo^ and TCF-1^Lo^ CD25^Hi^ antigen-specific cells distinguishable during the activation and early expansion phase, were not functionally exhausted, as evidenced by vigorous production of IFN-γ and TNF-α effector cytokine upon restimulation with cognate antigen, albeit the TCF1^Hi^ cells exhibited a modest trend towards decreased IFN-γ expression (Extended Data Fig 1C). Importantly, inverse association between the expression of CD25 and TCF-1 occurred similarly in TCR-transgenic as well as endogenous antigen-specific CD8 T cells of distinct specificities (D^b^Gp33- and D^b^GP276- specific tetramer+ CD8 T cells; Fig 1D Extended Fig 1A, 1D). Likewise, the precursor frequency of antigen-specific CD8 T cells also did not have a bearing on the generation of CD25^Lo^ TCF-1^Hi^ GzmB^Lo^ stem-like cells during chronic LCMV infection (Extended Data Fig 1A). While the kinetics of appearance of CD25^Lo^ TCF-1^Hi^ GzmB^Lo^ stem-like cells was slightly faster at higher precursor frequencies (Extended Data Fig 1A), the final CD8 T cell outcome during later stages of exhaustion was independent of their initial precursor frequencies (Extended Data Fig 2A-C). Collectively, these data demonstrate heterogeneity in early priming events during chronic viral infection, and establish an inverse association of IL-2Rα expression with TCF-1, and implicate a preferential enrichment of stem-like TCF-1^Hi^ cells in virus-specific CD8 T cells that express lower levels of IL-2Rα during priming.

### IL-2Ra^Lo^ cells are preferentially enriched for TCF-1^Hi^ cells, and give rise to long-lived TCF-1^Hi^ progenitor exhausted CD8 T cells during chronic infection

As shown in Fig 2A-B, CD25^Lo^ cells from GP33, GG276 and NP396-specific CD8 T cells all showed enrichment of TCF-1^Hi^ cells than CD25^Hi^ cells. We next sought to determine whether TCF-1^Hi^ CD25^Lo^ CD8 T cells identifiable during early stages of priming and expansion give rise to stem-like progenitor exhausted CD8 T cells during later stages of viral chronicity. We FACS-purified CD25^Lo^ and CD25^Hi^ D^b^GP33-specific CD8 T cells from day 3.5 LCMV_Cl-13_ infection, and tracked their long-term fate following adoptive transfer into infection-matched recipients (Fig 2C). Prior to adoptive transfer, The CD25^Lo^ subset of virus-specific CD8 T cells was preferentially enriched for TCF-1^Hi^ and Slamf6^Hi^ cells, whereas the CD25^Hi^ subset was enriched for GzmB^Hi^ and Tim-3^Hi^ cells (Extended Data Fig 2D). Compared to CD25^Lo^ subset, CD25^Hi^ cells also expressed higher levels of most inhibitory receptors analyzed (PD-1, Tim-3, Lag-3) (Extended Data Fig 2E), and key transcription factors implicated in regulating effector and T_Ex_ differentiation (Blimp-1, T-bet and Tox) (Extended Data Fig 2F). CD25^Hi^ cells also showed higher expression of cell proliferation markers (BrdU, Ki-67 and cMyc) (Extended Data Fig 2G), and were larger in size (Extended Data Fig 2H). These differences between CD25^Hi^ and CD25^Lo^ subsets were largely independent of their activation status, as both subsets showed potent upregulation of activation markers CD44 and CD69 and downregulation of CD127 and CD62L relative to naïve cells (Extended Data Fig 2H).

**FIGURE 2.**
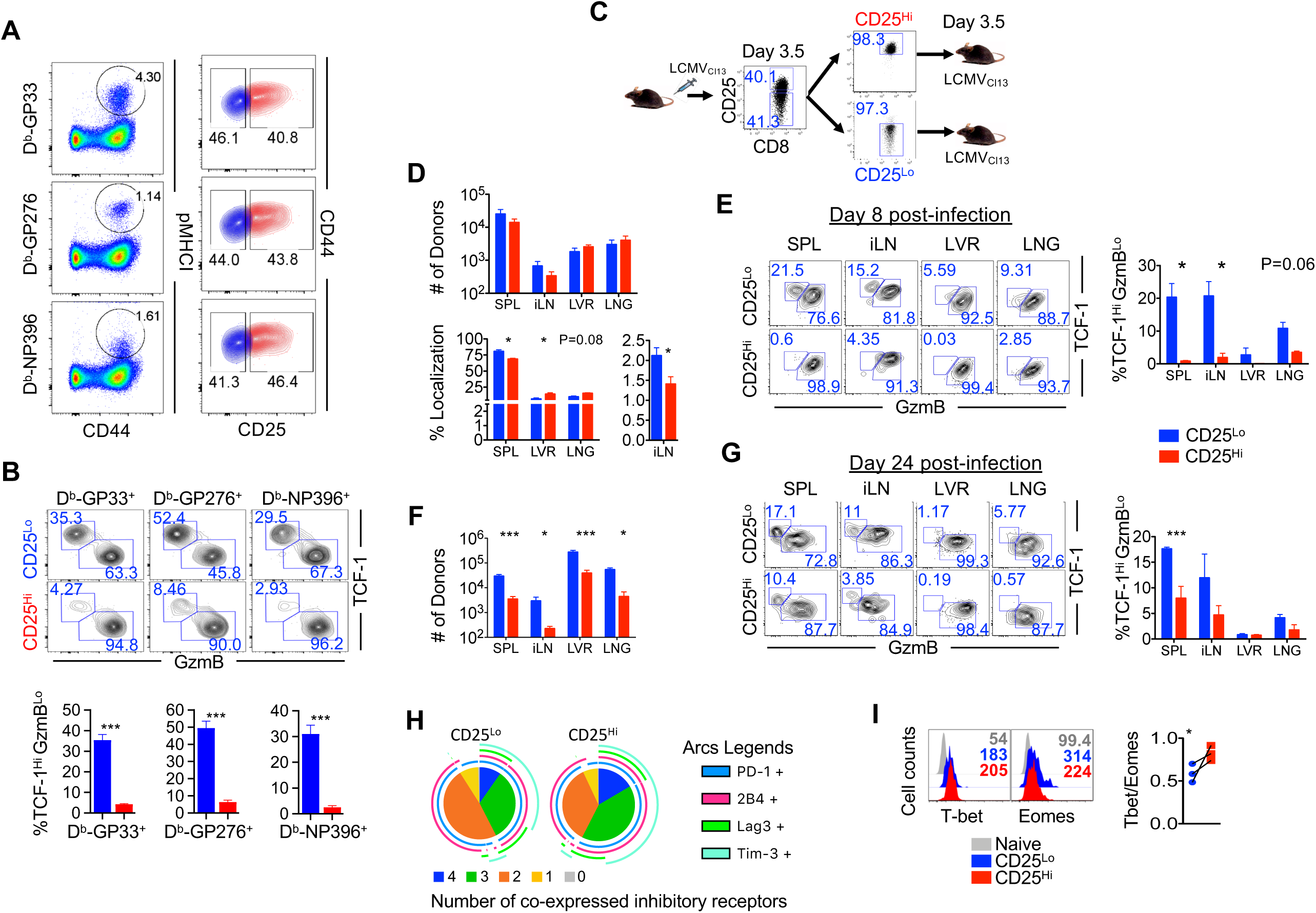
Stem-like TCF-1^Hi^ progenitor exhausted CD8 T cells that persist during chronic infection arise from CD25^Lo^ primed cells. **(A)** B6 mice infected with LCMV_Cl-13_ 5 days previously were assessed for CD25 expression on GP33, GP276 and NP396 specific CD8 T cells. (B) Tetramer+ CD8 T cells expressing low (blue) or high (red) CD25 expression were analyzed TCF-1 and GzmB expression. %TCF-1Hi GzmBLo stem-like CD8 T cells amongst CD25Lo and CD25Hi antigen-specific CD8 T cells **(C)** are presented as bar graphs. Experimental set-up to investigate the long-term exhausted CD8 T cell fate outcomes of CD25^Hi^ and CD25^Lo^ LCMV-specific CD8 T cells isolated during priming. WT P14 cells (1×10^6^) were adoptively transferred into naïve B6 recipient mice and then infected with LCMV_Cl-13_ 1 day later. CD25^Lo^ and CD25^Hi^ cells were FACS purified 3.5 days after LCMV_Cl-13_ infection. Congenically mismatched CD25^Hi^ or CD25^Lo^ donor cells (∼1×10^5^) were transferred into infection-matched C57Bl/6 mice. **(D)** Bar graphs show numbers and % localization of donor cells in spleen (SPL), inguinal lymph nodes (iLN), liver (LVR) and lung (LNG) at day 8 after infection. **(E)** Flow-cytometry plots of TCF-1 and GzmB co-expression in transferred CD25^Hi^ or CD25^Lo^ donor cells are presented from the indicated tissues, 8 days after infection. Representative flow data are shown along with a bar graph of % stem-like TCF-1^Hi^ GzmB^Lo^ CD8 T cells. **(F)** Bar graphs show numbers of transferred CD25^Hi^ and CD25^Lo^ P14 CD8 T donor cells in tissues on day 24 after LCMV_Cl13_ infection. **(G)** Flow-cytometry plots of TCF-1 and GzmB co-expression in transferred CD25^Hi^ or CD25^Lo^ donor cells are presented from the indicated tissues, 24 days after infection. Representative flow data are shown along with a bar graph of % stem-like TCF-1^Hi^ GzmB^Lo^ CD8 T cells. **(H)** SPICE (simplified presentation of incredibly complex evaluations) plots illustrating co-expression of the inhibitory receptors (IR) PD-1, Tim-3, LAG3, 2B4 on transferred CD25^Hi^ or CD25 ^Lo^ P14 CD8 T cells are presented. **(I)** Relative expression of T-bet and eomesodermin transcription factors in CD25^Lo^ and CD25^Hi^ donors are presented as histogram plots. Bar graphs depict the expression of T-bet and Eomes on FACS sorted and transferred CD25^Hi^ (Red) or CD25^Lo^ (Blue) P14 CD8 T cells at day 8 post-infection; grey histograms show endogenous CD44^Lo^ naïve CD8 T cells. Number represents MFI of respective marker. Bar graphs display mean and SEM. Data are representative of at least two independent experiments with n=5-10 mice per group. Unpaired Student t-test was used with statistical significance in difference of means represented as * (P ≤ 0.05), ** (P ≤ 0.01), *** (P ≤ 0.001).

Notwithstanding these initial differences, consistent with similar expression levels of pro-survival molecule Bcl-2 (Extended Data Fig 2G), the CD25^Hi^ and CD25^Lo^ purified donors engrafted similarly. Both donors also expanded to largely similar levels at the peak of CD8 T cell expansion (day 8 post-infection with LCMV_Cl-13_) (Fig 2D). The CD25^Lo^ donors (enriched in TCF-1^Hi^ Tim-3^Lo^ cells at the time of purification; Extended Data Fig 2I) maintained higher proportions of TCF-1^Hi^ Tim-3^Lo^ GzmB^Lo^ stem-like cells compared to CD25^Hi^ donors (Fig 2E), and preferentially localized to secondary lymphoid organs (spleen and lymph nodes) compared to peripheral lung and liver sites (Fig 2D). However, with progression of infection, there was a striking decline in the numbers of CD25^Hi^ donors, which largely disappeared, and were 10-20-fold lower in numbers than their CD25^Lo^ counterparts at day 24 post-infection in all tissues analyzed (Fig 2F). Greater recovery of CD25^Lo^ donors was associated with higher proportions of TCF-1^Hi^ GzmB^Lo^ (Fig 2G) CD62L^Hi^ Slamf6^Hi^ (Extended Data Fig 2J) stem-like cells, and lesser degree of exhaustion as evidenced by fewer donors co-expressing three or more inhibitory receptors (PD-1, Tim-3, Lag-3, 2B4) compared to the small surviving population of CD25^Hi^ donors (Fig 2H). In contrast, consistent with a more terminally exhausted phenotype, CD25^Hi^ donors also expressed higher levels of Tbet/Eomes ratio (Fig 2I, Extended Data Fig 2K). Notably, the CD25^Lo^ and CD25^Hi^ donor cells, minimally impacted the infectious milieu as demonstrated by largely similar endogenous D^b^GP33-specific CD8 T cell responses over time (Extended Data Fig 2L), and similar proportions of stem-like cells (Extended Data Fig 2M) and PD-1 expression (Extended Data Fig 2N) in endogenous virus-specific CD8 T cells from the recipients. Collectively, these data demonstrate that CD25^Lo^ TCF-1^Hi^ cells distinguishable during early stages of priming and expansion preferentially differentiate into stem-like progenitor exhausted CD8 T cells during later stages of viral chronicity.

### Stem-like TCF-1^Hi^ progenitor CD8 T cells arising from IL-2Rα^Lo^ precursors exhibit augmented responsiveness to PD-1 checkpoint blockade immunotherapy during exhaustion

Compared to TCF-1^Lo^ terminally exhausted CD8 T cells, TCF-1^Hi^ progenitor exhausted CD8 T cells mount superior expansion in response to PD-1 CBI in both chronic infection and tumor model studies ^3–5, 7, 9, 15^. Hence, we next assessed this key functional property of stem-like TCF-1^Hi^ progenitor cells arising from CD25^Lo^ precursors programmed during early stages of priming and expansion. As in Fig 2, CD25^Lo^ and CD25^Hi^ D^b^GP33-specific CD8 T cells were FACS-purified from day 3.5 LCMV_Cl-13_ infection, adoptively transferred into infection-matched recipient mice, and allowed to differentiate under conditions of viral chronicity. The recipient mice were then subjected to PD-1 checkpoint blockade immunotherapy, and donor cell expansion, phenotype and function were assessed (Fig 3A). Consistent with their stem-like less exhausted TCF-1^Hi^ Tim-3^Lo^ GzmB^Lo^ Slamf6^Hi^ CD62L^Hi^ phenotype (Extended Data Fig 2D-K), the CD25^Lo^ donors mounted significantly greater expansion in response to PD-1 CBI compared to CD25^Hi^ donors in all tissues analyzed (PBMC, spleen, lymph node, lung and liver) (Fig 3B, 3C, Extended Data Fig 3A), and better controlled viral loads (data not shown). However, minimal differences were observed in effector differentiation between CD25^Lo^ and CD25^Hi^ donors, as evidenced by largely similar levels of expression of GzmB, PD-1 and TNF-α following PD-1 CBI (Fig 3D, 3E). Notably, the responsiveness (expansion and effector differentiation) of endogenous D^b^GP33-specific CD8 T cells of recipient mice remained unaltered to PD-1 therapy, regardless of CD25^Lo^ or CD25^Hi^ donors being adoptively transferred into them (Extended Data Fig 3B, 3C), thus lending further support to the data in Extended Data Fig 2L-N showing that the adoptively transferred donor cells do not perturb the infectious milieu of recipient mice with respect to viral loads, immune factors, etc. Together, these data establish that functionally potent TCF-1^Hi^ stem-like progenitor cells – capable of vigorous expansion in response to anti-PD-1 therapy – develop from CD25^Lo^ precursors programmed during early stages of chronic viral infection. These observations prompted us to hypothesize that tempered IL-2 signals during T cell priming serve to program the long-lived self-renewing subset of stem-like exhausted CD8 T cells capable of robust responses to anti-PD-1 therapy.

**FIGURE 3.**
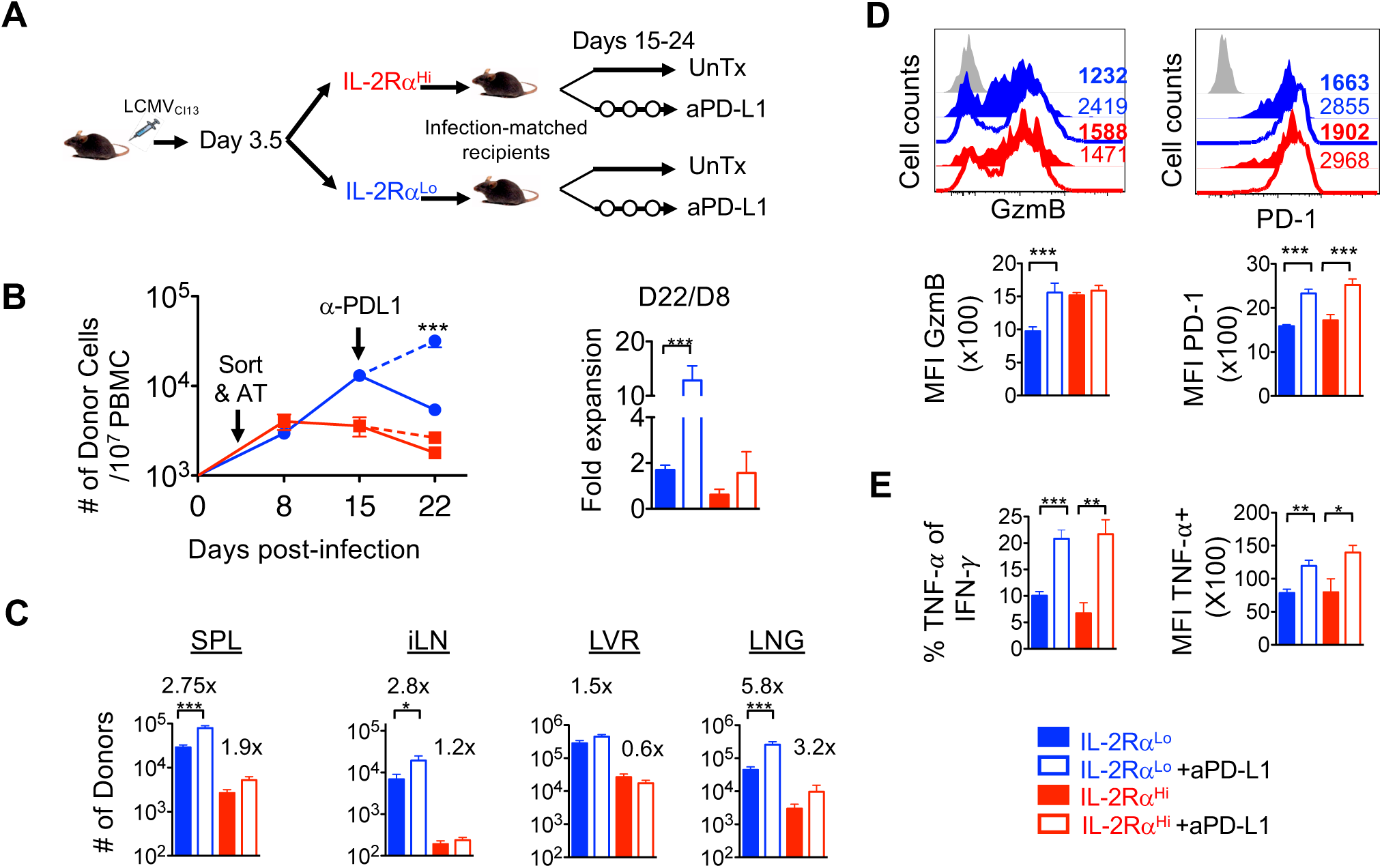
Stem-like TCF-1^Hi^ progenitor CD8 T cells arising from IL-2Rα precursors exhibit augmented responsiveness to PD-1 checkpoint blockade immunotherapy during exhaustion. **(A)** Experimental set-up. CD25^Lo^ and CD25^Hi^ cells were FACS purified at day 3.5 after infection, and adoptively transferred into infection-matched congenically distinct recipient mice (as in Fig 1). Mice were subsequently treated with anti-PD-L1 antibody (200 μg) every 3 days from day 15 to 24, and donor cells were assessed for expansion and effector function. **(B)** Longitudinal analysis of CD25^Lo^ and CD25^Hi^ donors in blood of chronically infected mice. Numbers of CD8^+^ Ly5.1^+^ donor cells per 10×10^6^ PBMC are depicted at indicated time points after infection. Bar graphs show fold expansion of donor cells at day 22 in blood after infection in treated and untreated groups receiving CD25^Lo^ or CD25^Hi^ donor cells. Data are representative of 2 independent experiments with n=4-6 mice per group. **(D)** Absolute numbers of donor cells in the presence or absence of anti-PD-L1 blockade therapy. Bar graphs show absolute numbers of donor cells in indicated tissues. Numbers above bars depict fold-change in numbers of donor cells after treatment with aPD-L1. **(D)** Expression of effector molecule granzyme B and inhibitory receptor PD-1 in CD25^Lo^ and CD25^Hi^ donors in response to anti-PD-L1 blockade therapy. Representative histograms and composite bar graphs show GzmB and PD-1 expression in donor CD8 T cells in the spleen at day 24 after infection. Numbers depict mean fluorescence intensity (MFI) of expression of respective markers. **(E)** Expression of effector cytokines by CD25^Lo^ and CD25^Hi^ donors in response to anti-PD-L1 blockade therapy. Bar graphs show % IFN-γ and TNF-α double positive donor cells, and MFI of TNF-α expression in CD25^Hi^ and CD25^Lo^ donor CD8 T cells from spleens following *in vitro* stimulation of splenocytes with GP33 peptide for 5 hrs. Bar graphs display mean, and SEM. Data are representative of at least 2 independent experiments with n=4-6 mice per group. One-way ANOVA with Tukey post-test was used to compare differences between groups. Statistical significance in difference of means is represented as * (P ≤ 0.05), ** (P ≤ 0.01), *** (P ≤ 0.001).

### Functional role of IL-2 signals in programming TCF-1^Hi^ stem-like precursors during chronic LCMV infection

Higher expression of IL-2Rα was associated with increased STAT-5 phosphorylation (Extended Data Fig 4A), increased expression of downstream metabolic regulator cMyc (Extended Data Fig 2G), and higher levels of glycolysis and respiration (as indicated by increased extracellular acidification rate and oxygen consumption rate in a Seahorse biochemical analysis; Extended Data Fig 4B-D). Hence, we next sought to directly query the implied role of differential IL-2 signals in programming the development of TCF-1^Hi^ and TCF-1^Lo^ CD8 T cells in chronic LCMV infection. Towards this goal, we either increased IL-2 signals through exogenous administration of IL-2, or tempered IL-2 signals through *in vivo* antibody blockade during early stages of chronic LCMV infection (days 0-3.5), and assessed the development of TCF-1^Hi^ GzmB^Lo^ stem-like CD8 T cells (Fig 4A). Appropriate IL-2 signal modulation was confirmed by assessing the levels of key IL-2 signaling-dependent proteins, CD25 and GzmB (Fig 4B). At these early stages of TCR-driven proliferation, all three groups – untreated, IL-2 supplemented and IL-2 blocked – exhibited similar levels of activation, as assessed by significant upregulation of CD44 compared to naïve cells (Extended Data Fig 4E); underwent significant proliferation compared to naïve cells as assessed by largely similar BrdU incorporation, and expanded to largely similar levels (Fig 4B, Extended Data Fig 4F). Additionally, in the short programming window of 3-5 days, we did not observe any significant alterations in Treg cells due to IL-2 manipulation (Fig 4C).

**FIGURE 4.**
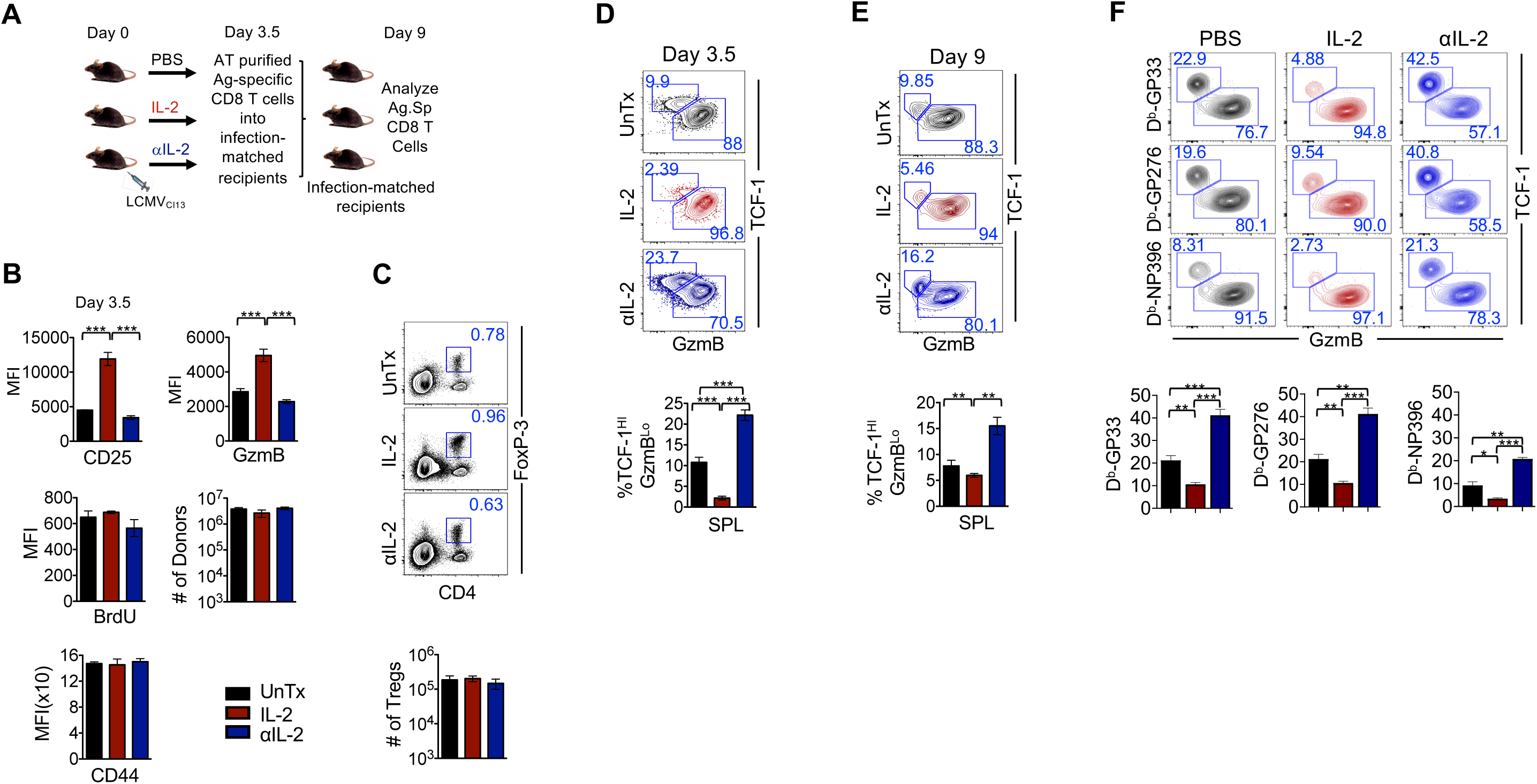
Attenuation of IL-2 signals in virus-specific CD8 T cells promotes the generation of TCF-1^Hi^ stem-like precursors during exhaustion. **(A)** Experimental setup for IL-2 enhancement or attenuation during T cell priming of chronic LCMV infection. Naïve C57Bl/6 mice adoptively transferred with 1×10^6^ P14 CD8 T cells were infected with LCMV_Cl-13_, and were treated intraperitoneally with PBS, high dose IL-2 (1.5 × 10^4^ IU) or anti-IL-2 complex from days 0-3.5 after infection. LCMV-specific CD8 T (4×10^5^) cells were isolated from splenocytes at day 3.5 post-infection, characterized, and transferred into infection-matched recipients. Donor cells were analyzed 6 days following transfer (Day 9 post-infection). **(B)** Activation, expansion and effector molecule expression under distinct IL-2 stimulatory conditions. Bar graphs show the MFI of expression of CD25, GzmB, BrdU, CD44 on donor cells in the untreated (UnTx) (Black), IL-2-treated (Red) and αIL-2 treated groups (Blue) at day 3.5 post-infection. Total numbers of donors at day 3.5 are also presented as bar graphs. **(C)** Representative flow-cytometry plots show the frequency of FoxP3+CD4 T regulatory cells (Treg) in the spleens of untreated or treated groups of mice at day 3.5 post-infection as indicated. Bar graph shows absolute numbers of Tregs in each group. **(D)** Assessment of stem-like precursors with IL-2 supplementation or blockade during T cell priming. Representative flow-cytometry plots of TCF-1 and GzmB co-expression on donor CD8 T cells, and composite bar graphs showing % TCF1^Hi^ GzmB^Lo^ stem-like precursors in the SPL at day 3.5 post-infection. Data are representative of 3-4 independent experiments with n=4-6 mice per group. **(E)** Persistence of stem-like CD8 T cell precursors primed under conditions of IL-2 attenuation. Representative flow-cytometry plots of TCF-1 and GzmB co-expression on donor CD8 T cells, and composite bar graphs showing % TCF1^Hi^ GzmB^Lo^ stem-like precursors in the SPL 6 days after adoptive transfer (∼9 days after LCMV_Cl-13_ infection) of donor cells primed under IL-2 supplementation or blockade conditions for 3.5 days. **(F)** C57Bl/6 mice infected with LCMV_Cl-13_ were treated withPBS, IL-2 or anti-IL2 for days 0-3.5. At day 8 post-infection, D^b^GP33, D^b^GP276 and D^b^NP396 tetramer+ CD8 T cells were assessed for expression of TCF-1 and GzmB. Representative raw data and Mean +/- SEM are presented as bar graphs. Paired Student t-test was used with statistical significance in difference of means represented as * (P ≤ 0.05), ** (P ≤ 0.01), *** (P ≤ 0.001). To compare differences between groups one-way ANOVA with Tukey post-test was used. Statistical significance in difference of means is represented as * (P ≤ 0.05), ** (P ≤ 0.01), *** (P ≤ 0.001).

As shown in Fig 4D, evident differences in the development of TCF-1^Hi^ antigen-specific CD8 T cells were noted with manipulation of IL-2 signals: whereas exogenous IL-2 administration from days 0-3.5 after LCMV_Cl-13_ infection led to reduced TCF-1^Hi^ GzmB^Lo^ cells, IL-2 blockade led to augmentation of TCF-1^Hi^ GzmB^Lo^ cells compared to untreated cells, in most secondary lymphoid and nonlymphoid tissues analyzed (spleen, lymph node, lung, liver) (Fig 4D, Extended Data Fig 4G). The donor cells primed in polarized high or low IL-2 conditions maintained their initial program, such that donors receiving increased IL-2 signals from days 0-3.5 remained lower for TCF-1^Hi^ GzmB^Lo^ cells even a week after adoptive transfer into infection-matched recipients with physiologic levels of IL-2 (Fig 4E). In contrast, day 3.5 donors programmed with tempered IL-2 signals remained higher for TCF-1^Hi^ GzmB^Lo^ cells following transfer into infection-matched recipients (Fig 4E). Increased IL-2 signaling specifically impaired the development of the TCF-1^Hi^ Tim-3^Lo^ stem-like subset, while mediating modest decrease in the intermediate TCF-1^Lo^ Tim-3^Lo^ subset, and a modest increase in the TCF-1^Lo^ Tim-3^Hi^ subset (Extended Data Fig 4H). Reduced IL-2 signaling showed the inverse outcome. Likewise, endogenous GP33-, GP276- and NP396-specific CD8 T cells also showed enhanced terminal differentiation upon IL-2 supplementation, and increased proportions of stem-like CD8 T cells upon IL-2 blockade in straight non-chimeric C57Bl/6 mice infectd with LCMVCl-13 and treated with IL-2 or anti-IL-2 from days 0-3 post-infection (Fig 4F). Notably, adoptive transfer of day 3.5 donors into day 3.5-infection matched recipients was conducted at low precursor frequencies, and did not alter the differentiation program of endogenous D^b^GP33-specific CD8 T cells, as indicated by similar numbers of endogenous D^b^GP33-specific CD8 T cells, which also contained largely similar levels of TCF-1^Hi^ GzmB^Lo^ cells regardless of receiving untreated, IL-2 treated or IL-2 blocked donors (Extended Data Fig 4I-J). These data establish a key functional role of tempered IL-2 signals in programming the development of TCF-1^Hi^ lineage in chronic viral infection.

### IL-2Rα is required in a T cell-intrinsic manner to drive terminal differentiation in chronic viral infection

Enrichment of TCF-1^Hi^ cells in the CD25^Lo^ subset of virus-specific CD8 T cells (Fig 2, Extended Data Fig 2), and the subsequent emergence of PD-1 checkpoint blockade immunotherapy responsive TCF-1^Hi^ stem-like progenitor cells from CD25^Lo^ precursors (Fig 3) further support the notion that IL-2 signal regulation might be acting at a T cell-specific level to program the generation of stem-like lineage, likely through physiologic modulation of CD25 expression. However, it is also plausible that IL-2 might drive the divergence of TCF-1^Hi^ and TCF-1^Lo^ CD8 T cell lineages through indirect effects on other immune cells such as effector CD4 T cells, Treg cells or APCs. Hence, we next engaged RNP-based Crispr/Cas9 methodology to specifically ablate CD25 expression in D^b^GP33-specific CD8 T cells from day 0 (Fig 5A-C, Extended Data Fig 5A-C) or day 2 after priming and activation (Fig 5D-F). We also used siRNA technology to transiently decrease CD25 expression between days 2-4 of priming and activation to query the timing of action of IL-2 signals in regulating the fate of TCF-1^Hi^ stem-like precursors (Extended Data Fig 5D-G). The development of TCF-1^Hi^ and TCF-1^Lo^ cells from wild-type (WT) and CD25 knockdown (KD) CD8 T cells was then assessed at day 8 after LCMV_Cl-13_ infection following transfer into naïve (in case of day 0 or siRNA-based transient knockdown) or day 2 infection-matched C57Bl/6 recipients (in case of day 2 knockdown) (Fig 5A, 5D). WT D^b^GP33-specific CD8 T cells were co-transferred with CD25 KD cells into the same recipient mice to ensure differentiation of both subsets in the same infectious milieu. The WT and KD donors were distinguishable from each other and from endogenous D^b^GP33-specific CD8 T cells using congenic Ly/Thy markers. The efficiency of transfection (Extended Data Fig 5A) and CD25 knockdown was confirmed by flow cytometry at days 2 (Fig 5B), 5 and 8 (Extended Data Fig 5b) after electroporation of guide RNAs. In the case of day 2 CD25 knockdown, CD25 expression levels were confirmed to be reduced at day 4 after activation as well as day 8 after infection (Fig 5E). In contrast to Crispr/Cas9-mediated deletion of *il2ra,* transient loss of CD25 was noted at day 2 after transfection in the case of siRNA-mediated knockdown, which was restored to WT levels at days 4 after knockdown and day 8 after infection (Extended Data Fig 5E).

**FIGURE 5.**
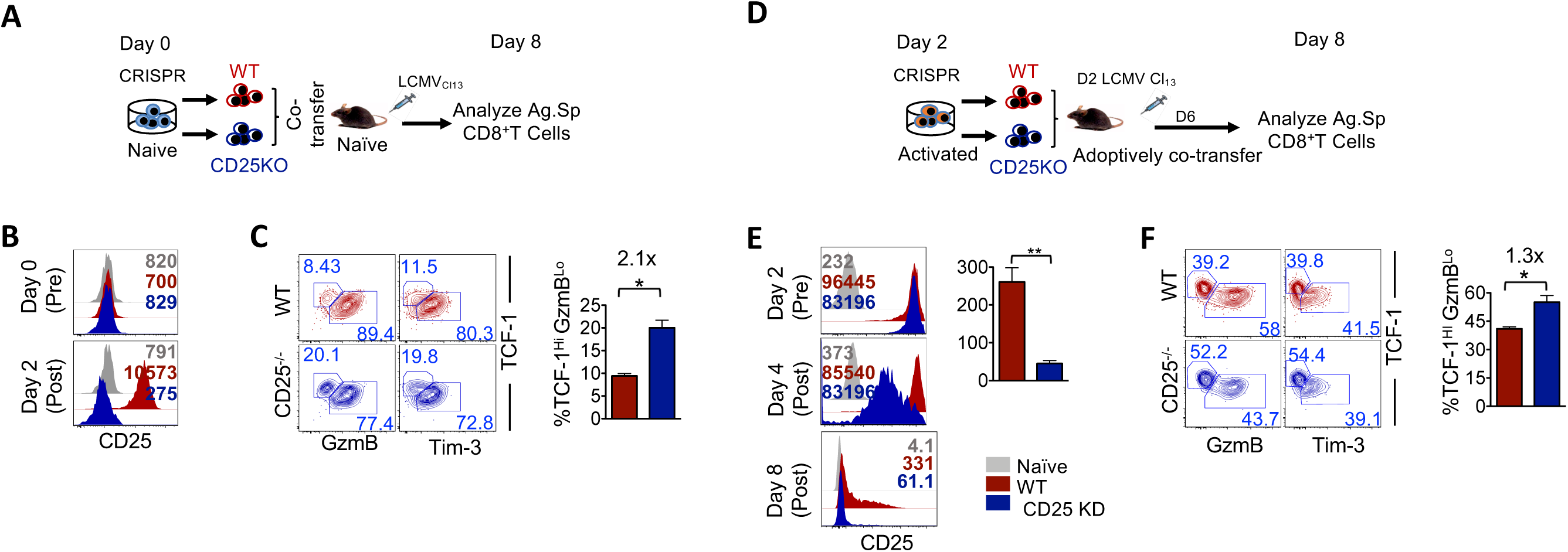
IL-2Rα exerts a T cell specific role in regulating the development of TCF-1^Hi^ stem-like CD8 T cells. **(A)** Experimental setup to query the CD8 T cell-intrinsic role of CD25 in programming TCF-1^Hi^ stem-like lineage from day 0. The gene encoding IL-2Rα was knocked out using RNP-based CRISPR-Cas9 technology. Naïve P14 CD8 T cells were electroporated with guide RNA targeting *il2ra* gene. Equal numbers of WT CD8 T cells or CD25 knockdown (KD) cells were adoptively co-transferred into naïve mice, which were then infected with LCMV_Cl-13_. Donor LCMV-specific CD8 T cells were analyzed after 8 days of infection. **(B)** The efficiency of CRISPR-Cas9-mediated deletion of *il2ra.* Representative histogram plots show the expression of CD25 in WT (Red), CD25 ^KD^ (Blue) and naïve cells (Gray) P14 CD8 T cells after 48hr of *in vitro* stimulation by plate bound anti-CD3 and anti-CD28. **(C)** Assessment of TCF1^Hi^ stem-like precursors in CD25 ablated CD8 T cells. Representative flow-cytometry plots of TCF-1 and GzmB or TCF-1 and Tim-3 co-expression on D0 ablated CD25 donor CD8 T cells in the spleen, along with composite bar graphs of % stem-like TCF-1^Hi^ GzmB^Lo^ CD8 T cells are presented. Data are representative of at least 2 independent repeats with n=3 mice per group (mean ± SEM). **(D)** Experimental setup to query the CD8 T cell-intrinsic role of CD25 in programming TCF-1^Hi^ stem-like lineage from day 2.. Day 2 activated P14 CD8 T cells were electroporated with guide RNA targeting *il2ra* gene. Equal numbers of WT CD8 T cells or CD25 KD cells were adoptively co-transferred into Day 2 chronically infected mice. Donor antigen specific CD8^+^ T cells were analyzed after 8 days of infection. **(E)** The efficiency of CRISPR-Cas9-mediated deletion of *il2ra* on D2 activated P14 CD8 T cells was confirmed by flow cytometry after 48hr of *in vitro* stimulation by plate bound anti-CD3 and anti-CD28. Histograms show the expression of CD25 in WT (Red), CD25 KD (Blue) and naïve cells (Gray). **(F)** Representative flow cytometry plots of TCF-1 and GzmB or TCF-1 and Tim-3 expression in D2 CD25-ablated donor CD8 T cells in spleens from day 8 LCMV_Cl-13_-infected mice are shown. Bar graph depicts % stem-like TCF-1^Hi^ GzmB^Lo^ CD8 T cells in gated WT and CD25 KD donor populations. To compare differences between groups Paired Student t-test or one-way ANOVA with Tukey post-test was used as appropriate, with statistical significance in difference of means represented as * (P ≤ 0.05), ** (P ≤ 0.01), *** (P ≤ 0.001).

In all three experimental conditions, we observed enhanced development of TCF-1^Hi^ GzmB^Lo^, Tim-3^Lo^ stem-like CD8 T cells upon CD25 knockdown, compared to wild-type CD8 T cells in spleen (Fig 5C, 5F, Extended Data Fig 5C, 5F). Even in the nonlymphoid liver site, where TCF-1^Hi^ cells are typically found at lower levels compared to lymphoid organs, we saw modest increase in TCF-1^Hi^ cells upon CD25 knockdown from day 0 (Extended Data Fig 5C) or transiently from days 0-4 after stimulation (Extended Data Fig 5F). As is typical for exhausted cells ^23–26^, Tox was expressed at higher levels than naïve cells, albeit CD25 ablation was associated with modest increase in Tox expression (Extended Data Fig 5G). Ablation of CD25 from day 0 led to a greater enhancement of TCF-1^Hi^ GzmB^Lo^, Tim-3^Lo^ stem-like progenitor exhausted CD8 T cells, compared to ablation of CD25 from day 2 or between days 0-4, thus suggesting that CD25 continues to exert an impact on the development of TCF-1^Hi^ cells at least up to 4 days after infection. These data establish that CD25 serves as a T cell-specific physiologic regulator of TCF-1^Hi^ stem-like cell development during early stages of chronic viral infection by modulating IL-2 signals.

### Late priming of CD8 T cells gives rise to TCF-1^Hi^ stem-like precursor CD8 T cells

In viral infections, IL-2 levels in the spleen typically peak around day 3 after infection and progressively decrease to baseline levels by day 6^38, 39^ (Extended Data Fig 6A). Hence, we hypothesized that T cell priming during later stages of expansion, when IL-2 levels are declining following chronic LCMV infection, may promote the development of TCF-1^Hi^ stem-like CD8 T cells. To test this, we adoptively transferred naïve D^b^GP33-specific CD8 T cells at day 0, 1, 2 or 3 after infection, and analyzed the level of TCF-1^Hi^ GzmB^Lo^ Slamf6^Hi^ stem-like CD8 T cells in the secondary lymphoid (spleen and lymph node) and nonlymphoid tissues (liver and lung) at day 8.5 LCMV_Cl-13_ after infection (Fig 6). We noted that conditions of late priming and curtailed duration of stimulation (days 3-8.5) were more conducive to the development of stem-like CD8 T cells in all tissues analyzed, compared to prolonged stimulation from days 0-8.5 after infection (Fig 6A-B, Extended Data Fig 6B). Curtailed stimulation during later stages from days 3-8.5 was also associated with lesser proliferation (as indicated by Ki-67 staining), lower level of Tim-3 expression and largely similar levels of PD-1 expression (Extended Data Fig 6C). Likewise, curtailing the duration of stimulation to early stages of infection (days 0-3.5 and days 0-5.5) induced higher levels of TCF-1^Hi^ GzmB^Lo^ stem-like virus-specific CD8 T cells, compared to longer durations of stimulation from days 0-6 or days 0-8 (Extended Data Fig 6D-E). However, the proportions of stem-like CD8 T cells were lower when cells were stimulated during early stages of infection (when IL-2 levels are higher), than during later stages of priming when IL-2 levels decline (Extended Data Figure 6B, 6E). Early stimulation from days 0-5.5 induced about 10-15% TCF-1^Hi^ cells, compared to stimulation during later stages of expansion from days 3-8.5 stimulation, which induced about 40% TCF-1^Hi^ cells despite identical duration of stimulation (Fig 6A, Extended Data Fig 6B). These data indicate that shorter duration of stimulation and later priming are more conducive for the development of TCF-1^Hi^ stem-like CD8 T cells.

**FIGURE 6.**
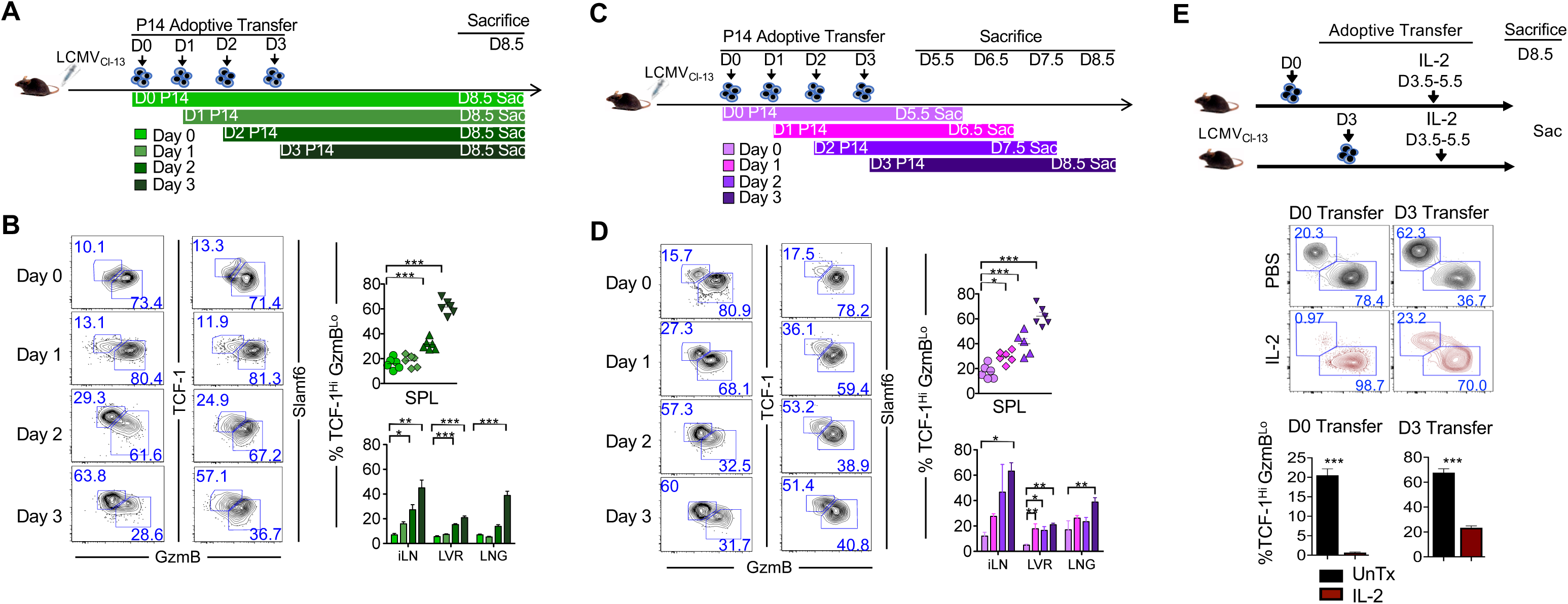
Late priming of CD8 T cells gives rise to TCF-1^Hi^ stem-like precursor CD8 T cells. **(A)** Experimental setup. LCMV_Cl13_-infected C57Bl/6 mice were adoptively transferred with 2.5×10^3^ of WT P14 CD8 T cells at day 0, 1, 2 or 3 after infection. Donor CD8 T cells were analyzed 8.5 days after infection. **(B)** Representative flow cytometry plots of TCF-1 and GzmB or TCF-1 and Slamf6 expression in donor CD8 T cells at day 8.5 post-infection are presented from spleen. Bar graphs depict % TCF-1^Hi^ GzmB^Lo^ CD8 T cells in the SPL (top) and iLN, LVR and LNG (Bottom) at 8.5 days after infection. **(C)** Experimental setup. LCMV_Cl-13_ infected C57Bl/6 mice were adoptively transferred with 2.5×10^3^ of WT P14 at day 0, 1, 2 and 3 after infection. Donor CD8 T cells were analyzed 5.5 days after P14 transfer. **(D)** Characterization of stem-like CD8 T cell fates upon early versus late priming in LCMV_Cl-13_ infection. Representative flow cytometry plots of TCF-1 and GzmB or TCF-1 and Slamf6 co-expression on donor CD8 T at indicated time-points (days 5.5, 6.5, 7.5 or 8.5) after infection are presented. Bar graphs depict % stem-like TCF-1^Hi^ GzmB^Lo^ CD8 T cells in the SPL (top), or iLN, LVR and LNG (Bottom) at 5.5 days after P14 transfer. **(E)** Low dose P14 CD8 T cells were adoptively transferred into B6 mice at D0 or D3 after LCMV_Cl-13_ infection, which were then treated with PBS or IL-2. P14 donor CD8 T cells were analyzed for proportions of TCF-1 and GzmB to assess the proportions of stem-like and terminally differentiated lineages.

To directly investigate whether priming during later stages of chronic LCMV infection better supports the development of TCF-1^Hi^ stem-like CD8 T cells compared to earlier stages, we next scanned how a 5.5 day window of stimulation from days 0-5.5, 1-6.5, 2-7.5 or 3-8.5 impacted the development of stem-like CD8 T cells (Fig 6C). Thus, maintaining the duration of stimulation the same (5.5 days), we controlled the timing of stimulation by adoptively transferring naïve D^b^GP33-specific CD8 T cells at day 0, 1, 2 or 3 after infection, and analyzed the level of TCF-1^Hi^ GzmB^Lo^ Slamf6^Hi^ stem-like CD8 T cells. Stem-like CD8 T cells were maximally induced in the days 3-8.5 stimulation window in all secondary lymphoid (spleen and lymph node) and nonlymphoid tissues (liver and lung) analyzed (Fig 6D, Extended Data Fig 6F). We noted a progressive increase in TCF-1^Hi^ GzmB^Lo^ Slamf6^Hi^ stem-like CD8 T cells with later priming, such that donor cells transferred at day 3 after infection exhibited the highest proportions of stem-like antigen-specific CD8 T cells, whereas cells primed during early stages of infection (days 0-5.5) exhibited the least amounts (Fig 6D). No significant differences in proliferation, or expression of canonical inhibitory receptors Tim-3 and PD-1 was noted between groups, consistent with largely similar duration of antigenic stimulation (Extended Data Fig 6G). Notably, IL-2 supplementation from days 3-8.5 reduced the proportions of TCF-1^Hi^ cells in both day 0 and day 3 spiked in cells (Fig 6E). Likewise, endogenous GP33, GP276 and NP396 CD8 T cells also showed reduced development of TCF-1^Hi^ cells following treatment with IL-2 from days 3-8.5 (Extended Data Fig 6H). Collectively, these data support the notion that differential IL-2 signals in these early versus late priming windows may be the underlying cause. These findings implicate differential priming of antigen-specific CD8 T cells during early versus late stages of expansion in chronic LCMV infection, and further demonstrate that longer duration of stimulation, and stimulation during early stages of infection in higher IL-2 conditions are detrimental to the development of TCF-1^Hi^ stem-like CD8 T cells. In contrast, shorter duration of stimulation, during later stages of infection with lower IL-2 levels, are conducive for the development of stem-like TCF1^Hi^ CD8 T cells during exhaustion.

### Correlations of IL-2 signaling signature with stem-like lineage and checkpoint blockade immunotherapy responses in melanoma, head and neck cancer and lung cancer patients

Analogous to chronic viral infections, functional exhaustion of tumor-reactive CD8 T cells is well established in solid tumors ^1^. Importantly, recent studies have identified a wide spectrum of heterogeneity in exhausted CD8 T cell pool, ranging from stem-like to terminally exhausted tumor-reactive CD8 T cells in a variety of patient tumor samples including melanoma, lung carcinoma and head and neck cancer ^7, 16–21^. In order to explore a potential relationship between IL-2 signaling and T cell exhaustion in patient solid tumor samples, we analyzed scRNA-seq data from melanoma, HPV+ head and neck cancer and lung carcinoma samples ^7, 16, 21^. As in the case of LCMV-specific CD8 T cells, tumor infiltrating lymphocytes (TIL) from patient melanoma samples showed an inverse association between TCF-1 and IL-2 signaling signature genes in uniform manifold approximation and projection (UMAP) analyses of scRNA-seq data (Fig 7A). TILs expressing lower levels of gene encoding TCF-1 exhibited higher expression of Granzyme B, Tim-3 and PD-1 encoding genes, alongside increased IL-2 signaling signature genes such as STAT5 targets and Blimp-1 (encoded by *PRDM1*) downstream of IL-2 (Fig 7A). In contrast, higher TCF-1 gene expression was directly related to higher CD62L and reduced IL-2 signaling signature genes (Fig 7A) as seen in LCMV-specific CD8 T cells. Unsupervised clustering of CD8 T cells into stem-like, transitory effector-like and terminally exhausted subsets based on gene expression profiles (Extended Data Fig 7A-B) (as validated by relative enrichment of memory or exhaustion signatures; Fig 7B, Extended Data Fig 7C) further confirmed a preferential enrichment of IL-2 signaling signature in the more differentiated transitory effector and terminally exhausted subsets compared to the less-differentiated stem-like cells (Fig 7C). Importantly, IL-2 signaling signature was also found to be enriched in TILs of melanoma patients who did not benefit from checkpoint blockade (non-responders), whereas patients with robust responses to therapy (responders) showed significantly lower IL-2 signaling signature enrichment scores (Fig 4D, Extended Data Fig 7D). Thus, these data link IL-2 signaling with increased TIL exhaustion and poor responsiveness to PD-1 CBI in melanoma patients.

**FIGURE 7.**
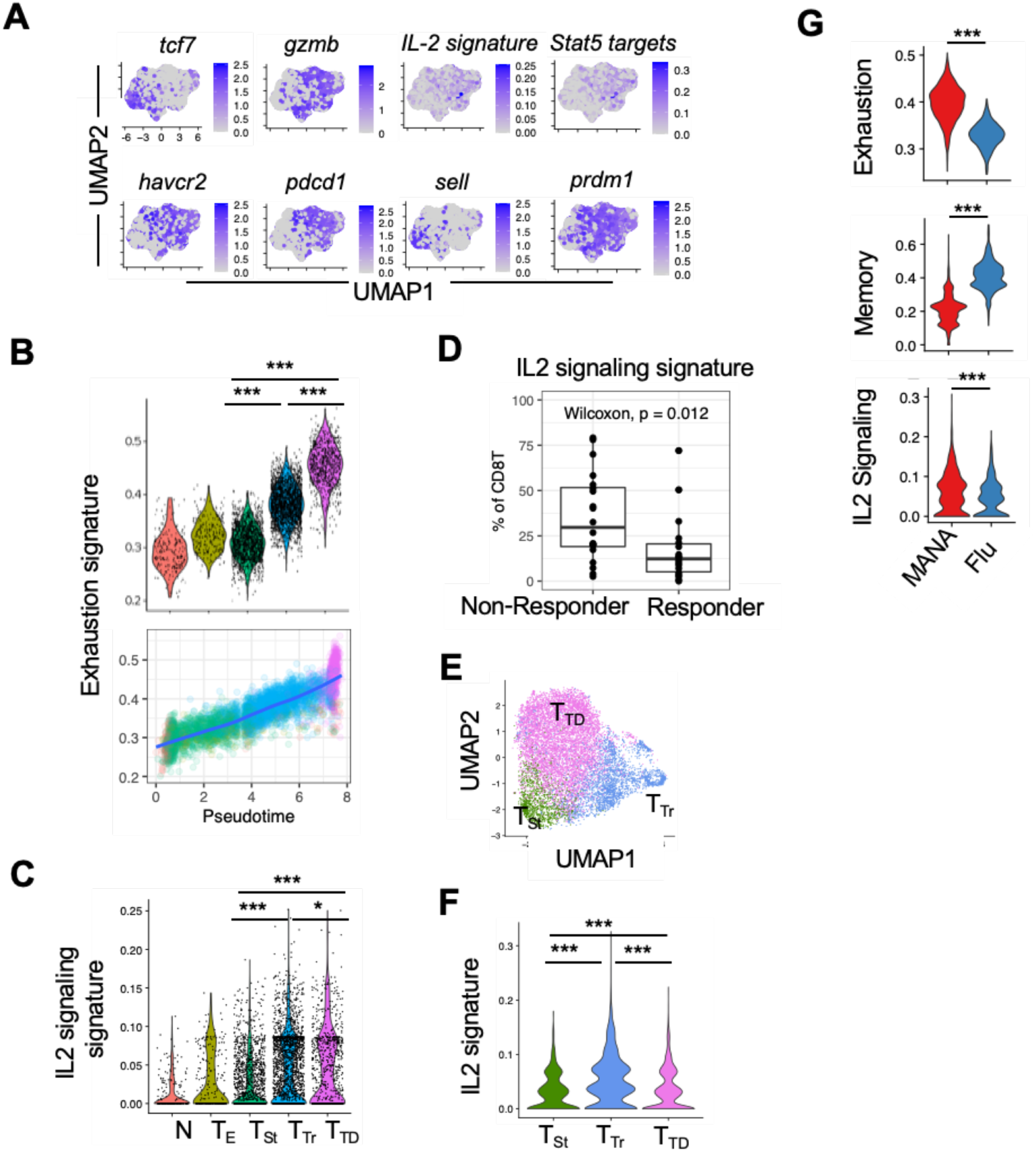
IL-2 signaling signatures are inversely associated with TCF-1^Hi^ lineage and anti-PD-1 responses in melanoma, HPV+ head and neck cancer and lung cancer patients. **(A)** Expression of key exhaustion and memory genes and IL2 signaling score in CD8 T cells in melanoma tumors from checkpoint blockade immunotherapy treated patient dataset from Sade-Feldman et al., 2018 UMAP plots of scRNA-seq of CD8 T cells from 48 melanoma tumors for activation/exhaustion associated genes (*PDCD1, HAVCR2, GZMB*), stem cell associated genes (*TCF7, SELL*) or a composite IL2 signature (FUNG_IL2_SIGNALING_1). Genes containing STAT5 binding sites and Blimp-1 (encoded by *PRDM*) downstream targets of IL-2 are also presented as UMAP plots. Genes downstream of IL2 signaling that also contain STAT5 binding sites (FUNG_IL2_TARGETS_WITH_STAT5_BINDING_SITES_T1) were used to calculate the STAT5 Target Binding score. (Transcript levels are color-coded: grey, not expressed; purple, expressed). **(B)** Exhaustion signature scores are shown for indicated CD8 T cell clusters from melanoma tumors and across pseudotime. **(C)** Signature of genes from the FUNG_IL2_SIGNALING_1 dataset was created and scored across the Naïve (N), Effector (E), Stem-like (T_St_) exhausted, Trasitory-exhausted (T_Tr_), and Terminally differentiated exhausted (T_TD_) clusters. **(D)** Association between immunotherapy response and IL2 signature scores in CD8 T cells. The frequency of CD8 T cells in either of the clusters with high IL2-signatures (T_Tr_ and T_TD_) as a portion of all CD8 T cells were compared between patients who responded or did not respond to checkpoint blockade. Wilcoxon ranked-sum test used to establish statistical significance. **(E)** Phenotypes of HPV- specific CD8 T cells in HPV driven tumors. CD8 T cells binding to MHC-I tetramers containing HPV peptides were sorted using flow cytometry and scRNA-seq was performed on the sorted cells from 12 patients. Unbiased clustering and UMAP plots were generated as described in the original publication ^16^. Unbiased clusters were labeled as stem-like (T_St_), transitory (T_Tr_) or terminally differentiated (T_TD_) based on their expression of memory and exhaustion genes and concordance with the original publication. **(F)** IL2 signature score in exhausted CD8 T cell clusters identified in panel d from HPV+ head and neck tumors. A signature score of genes from the FUNG_IL2_SIGNALING_1 dataset was created and scored across the clusters. Statistical significance was assigned using Wilcoxon ranked sum tests. **(G)** Violin plots of exhaustion, memory and IL2 signaling related gene signatures in neoantigen and influenza-specific CD8 T cells from NSCLC samples. CD3 T cells from 15 non-small cell lung cancer (NSCLC)-tumors were isolated and single cell RNA-seq was performed on these cells. In parallel the MANAFEST and viraFEST assays were performed to identify mutation associated neoantigens (MANA) or influenza specific T cell receptors. Cancer-specific signature neoantigen score was derived from ^55^. CD8 T cells with either MANA or influenza-specific T cells were isolated *in silico* and UMAP plots of these cells were performed on dataset from ^21^. Signature scores were calculated using expression of exhaustion- or memory -associated genes and genes downstream of IL2 signaling (FUNG_IL2_SIGNALING_1). Wilcoxon ranked-sum tests were used to assign statistical significance. Statistical significance represented as * (P ≤ 0.05), ** (P ≤ 0.01), *** (P ≤ 0.001).

In virally-associated human papillomavirus (HPV)+ head and neck cancers as well, UMAP analysis of scRNA-seq data from HPV tetramer+ TILs showed inverse expression patterns of TCF-1 with GzmB, Tim-3, PD-1 and IL-2 signaling, whereas higher TCF-1 expression correlated with higher CD62L expression, as in case of melanoma (Extended Data Fig 7E). Importantly, we noted an evident enrichment of IL-2 signaling signature in terminally differentiated antigen-specific TIL subsets compared to the stem-like cells defined by unsupervised cluster analysis (Fig 7E-F; Extended Data Fig 7F-G). Consistent with higher IL-2 signaling in more differentiated HPV+ TILs, we noted increased expression of STAT5 targets downstream of IL-2 and Blimp-1 (Extended Data Fig 7H). Likewise, scRNA-seq data analysis of neoantigen-specific and influenza A virus-specific TILs from lung cancer patients (Extended Data Fig 7I-J) also showed preferential enrichment of exhaustion and IL-2 signaling signatures (Fig 7G) (including STAT5 and Blimp-1, Extended Data Fig 7K) compared to influenza A virus-specific T cells from the same environment, which were more memory-like (Fig 7G). Collectively, these studies show a direct regulatory effect of IL-2 signaling in the development of stem-like and terminally exhausted CD8 T cell lineages and checkpoint blockade immunotherapy responses during chronic viral infection and solid tumors.

## DISCUSSION

IL-2 – a key immunomodulatory cytokine, capable of effecting both immunostimulatory and immunosuppressive physiologic outcomes ^39^ – is shown here to act in a rheostatic manner to program the development of stem-like TCF-1^Hi^ exhausted CD8 T cell precursors at low signaling intensity, but drive terminal exhaustion under strong/prolonged signaling conditions. Our findings of *in vivo* fate programming of stem-like exhausted CD8 T cells by attenuated IL-2 signals in chronic viral infection, and direct association of increased IL-2 signaling with more terminal differentiation in intra-tumoral tumor-reactive T cells from patients with a variety of solid tumors such as melanoma, lung cancer and virally-induced head and neck cancer have implications in two main areas: first, for PD-1-based therapy of patients with chronic viral infections and solid tumors; and second for adoptive T cell therapeutic strategies for chronic viral infections and cancers.

The efficacy of PD-1 based therapies is critically dependent on the presence of TCF-1^Hi^ stem-like exhausted CD8 T cells, capable of vigorous expansion and differentiation into effector CTLs for expeditious and efficacious control of virus or tumors ^3–5, 7, 9, 15^. Given that new thymic emigrants continue to be recruited into the virus- or cancer-specific immune response even in established disease ^4, 40^, our studies present regulated IL-2 signaling as a promising new preconditioning strategy that may be leveraged to improve clinical outcomes of PD-1-based therapies by shifting the balance in favor of stem-like exhausted CD8 T cell differentiation in ongoing cancers and chronic infections. This strategy to precondition an augmented response to PD-1-based therapies by IL-2 signal attenuation and augmentation of stem-like T cell subset prior to initiation of PD-1 checkpoint blockade is distinct from combination treatment with IL-2 and PD-1 checkpoint blockade, which is designed to boost effector responses following release of PD-1 brakes ^41^. Additionally, mechanisms that skew the balance of dendritic cells in favor of IL-2 non-producing subsets in the tumor microenvironment or during chronic infections may also be beneficial for augmenting T cell stemness and responsiveness to checkpoint blockade immunotherapy.

In the context of ACT, IL-2 has been classically used for expansion of therapeutic lymphocytes *in vitro* and for boosting their effector functions in patients following adoptive transfer ^42^. *In vivo* IL-2 administration offers beneficial effects by boosting effector responses, and several IL-2 biologics with targeted effects on anti-tumor T cells (and not Treg cells) are being evaluated ^42–44^. Our studies show that strong/prolonged IL-2 signaling eventually drives terminal exhaustion, whereas tempered IL-2 signals promote the generation of long-lived, multipotent stem-like antigen-specific T cells. These findings support the notion that tempering IL-2 signals during therapeutic T cell product generation will be beneficial for mediating long-term tumor control and immunosurveillance post-ACT by enhancing stem-like cells in the therapeutic product even in the controlled *in vitro* setting of T cell stimulation ^45^. Moreover, our studies present IL-2Rα and other components of the IL-2 signaling axis such as IL-2Rβ ^46^, Blimp-1 ^47^ and T cell-derived IL-2 ^48, 49^ as putative genetic engineering targets for enhancing the stemness and long-term therapeutic efficacy of TCR- or CAR-modified T cell monotherapy. This strategy is also expected to promote responsiveness of therapeutic product to subsequent PD-1 checkpoint blockade. Thus, our studies have broad implications in improving clinical outcomes of anti-PD-1 and ACT therapies for chronic infections and cancers by providing a unique preconditioning strategy of boosting long-lived, therapy-responsive stem-like CD8 T cells through IL-2 signal attenuation.

## METHODS

### Mice and infection

Four week old C57BL/6 mice were purchased from the Jackson Laboratory (Bar Harbor, ME, USA). Thy1.1+ and Ly5.1+ H-2D^b^GP33-specific TCR-transgenic P14 mice, fully backcrossed onto the C57BL/6 background, were maintained in our colony. IL-2-Cre^+^ mice were crossed with Rosa26 TdTomato reporter mice and bred in house. In all experiments, sex- and age-matched mice were used. All animals were used in accordance with SCRI Institutional Animal Care and Use Committee guidelines. LCMV Clone 13 strain (LCMV_Cl-13_) was propagated, titered, and used for infections as previously^50^. Mice were injected intravenously (i.v.) with LCMV_Cl13_ (2×10^6^ PFU) 12-16 hrs following adoptive transfer of P14 cells.

### Adoptive T cell transfer

To generate P14 chimeric mice, naïve C57Bl/6 mice were adoptively transferred with 2.5 ×10^3^ (Low dose) or 1×10^6^ (High dose) WT D^b^GP33-specific P14 CD8 T cells 12-16 hrs prior to infection with LCMV_Cl13_. Early T cell priming and expansion were analyzed at Day 3.5 in case of high dose P14 chimeras, and at day 5.5 for low dose P14 chimeric mice. FACS-purified CD25^Hi^ or CD25^Lo^ CD8 T cells isolated at day 3.5 after infection were adoptively transferred into infection-matched recipients, at similar numbers (1×10^5^-5×10^5^). In the CD25 ablation experiments, naïve P14 cells that were permanently knocked down for CD25 expression (D0 CD25 KD) using Crispr/Cas9 or transiently ablated for CD25 expression using siRNA (Transient 25 KD) were co-transferred along with their respective WT P14 controls (∼2×10^3^) at 1:1 ratio into naïve recipient mice one day prior to LCMV_Cl-13_ infection. Day 2 activated P14 CD8 T cells ablated for CD25 expression (D2 CD25 KD) and WT P14 T cells were similarly co-transferred into infection matched recipient mice at 1:1 ratio of 200×10^3^ cells of each type.

### Flow cytometry and cell sorting

All antibodies were purchased from Biolegend (San Diego, CA, USA) except for TCF-1 (Cell signaling) and Slamf6 (BD Biosciences). MHC class I tetramers were made as described on the NIH tetramer core facility protocol and used as previously ^51^. Single-cell suspensions from spleens, inguinal lymph nodes, lungs, livers or PBMCs from mice were prepared as previously ^51–54^. All samples were resuspended in FACs buffer (1xPBS containing 1% FBS and 0.05% sodium azide) and stained with a fixable dead cell dye (LIVE/ DEAD zombie) along with extracellular fluorophore-conjugated antibodies to detect cell surface markers. For analysis of intracellular cytokines, splenocytes (2×10^6^) were stimulated *in vitro* with 0.2μg/mL GP33-41 peptide in the presence of Brefeldin A for 5h, followed by surface staining for CD8, Ly5.1, Thy1.1, and Thy1.2. Cells were then washed and stained for IFN-γ, TNF-α and IL-2 using the BD Fixation/permeabilization kit (BD Bioscience). Staining of intranuclear transcription factors was performed using eBiosciences Foxp3/Transcription Factor Staining protocol. Phosphostaining was performed on splenocytes direct *ex vivo* or following *in vitro* stimulation with IL-2 in complete RPMI 1640 containing 10% FBS and β-mercaptoethanol (cRPMI) at 37°C in 5% CO2. Cells were immediately fixed with 1.6% formaldehyde for 10 min followed by permeabilization in 80% ice-cold methanol for 30 min at 4°C. After rehydration in FACs buffer cells were stained with anti-pSTAT5 on ice for 45 min as we have done previously ^51^. For *in vivo* BrdU incorporation assay, mice were injected with 500μL BrdU (2mg/ml) intraperitoneally 12h prior to analysis using a BrdU flow kit (BD Biosciences) as previously ^52, 53^. Sample acquisition was performed on LSRII Fortessa (BD Biosciences, San Jose, CA). Data analysis was performed with FlowJo v. 9.9 software.

For cell sorting of CD8 T cells, cells were enriched from total splenocytes using the EasySep Mouse CD8+ T cell Isolation Kit (Biolegend) at day 3.5 post infection. The enriched CD8 T cells were then FACS-sorted into CD25^Hi^ and CD25^Lo^ cells, following gating of Thy1.2-CD44+ P14 population. Cells were sorted on JAZZ cell sorter (BD Bioscience) using a 70-micron nozzle.

### In vivo treatment

For PD-1 checkpoint blockade immunotherapy studies, CD25^Hi^ and CD25^Lo^ recipient mice were treated intraperitoneally (I.P.) with PBS containing anti-PD-L1 monoclonal antibody (200µg/mouse, clone 10F.9G2, BioXcell) as previously^51^. PDL-1 blockade was performed every three days starting from day 16 to day 25 post infection for a total of three injections. To regulate IL-2 signals during priming (Day 0 to Day 3.5 post infection), mice were treated with high dose hIL-2 (15,000 IU, administered twice-daily through intraperitoneal route). IL-2 signaling blockade was performed using a combination of JES61A12 and S4B6 anti-IL-2 antibodies at 200μg/ml (BioXCell) injected once daily through intraperitoneal route. Control mice received sterile 1X PBS.

### T cell activation and electroporation

CD8+ T cells were enriched from total splenocytes using the EasySep Mouse CD8+ T cell Isolation kit (Stem Cell). For permanent or transient CD25 ablation experiments at day 0, 4-6 x 10^6^ P14 CD8 T cells were plated in 6 well plates and pre-cultured in IL-7 (10ng/ml, PeproTech) at 37°C, 5% CO2 for 24 hrs prior to electroporation. For ablation of CD25 at day 2 post-activation, purified naïve P14 CD8 T cells were plated at 3×10^6^ cells/well in αCD3/αCD28 coated 6 well plates. Cells were pre-activated for 48h at 37°C, 5% CO2 prior to electroporation. Note that the preparation of CRISPR/Cas9 reagents was conducted based on guidance provided by IDT Alt-R CRISPR-Cas9 system. Naïve cells were resuspended in T buffer (Invitrogen MPK1096) and electroporated using 2200V, 10ms, 3 pulses, while activated CD8 T cells were resuspended in R buffer (Invitrogen MPK1096) and electroporated using 1600V, 10ms, 3 pulses. Following Neon transfection P14 CD8 T cells were transferred into warm cRPMI. Cells were rested at 37°C, 5% CO2 for 20-30min. The electroporation efficiency was evaluated for ATTO uptake by flow cytometry. Naïve and effector CD8 T cells were then plated on αCD3/αCD28 coated plates for activation and stained for CD25 expression 48h and 96h post electroporation.

### Seahorse Assay

Oxygen consumption (OCR) and extracellular acidification rates (ECAR) were measured using the XFe96 well Seahorse Analyzer (Agilent). Day 3.5 Sorted CD25^Hi^ and CD25^Lo^ were adhered onto Seahorse cell culture plates using poly-L-lysine (Sigma) at 1.5×10^5^ cells per well. Agilent Seahorse XF Cell Mito Stress Test Kit was used to perform Mitostress test in the presence of 10mM glucose (Agilent) with addition of Oligomycin, FCCP, Rotenone/Antimycin A and finally 2-Deoxyglucose (20mM) to obtain the zero-point value of extracellular acidification.

### RNA-seq Data analyses

#### Mouse samples

scRNA-seq (10x Genomics) data (<u>GSE119943</u>) from splenocytes isolated from P14 chimeric mice (adoptively transferred with 5000 D^b^GP33-specific P14 CD8 T cells) at day 4.5 after infection with LCMV_Cl-13_ ^36^ were downloaded and analyzed using Seurat (Version 2). Briefly, cells with percentage of mitochondrial genes below 0.05% were included. Cells with highest numbers of detected genes (top 0.2%) or lowest numbers of detected genes (bottom 0.2%) were considered as outliers and excluded from downstream analyses. Raw UMI counts were normalized to UMI count per million total counts and log-transformed. Variable genes were selected based on average expression and dispersion. Principal component analysis (PCA) analysis was performed using variable genes. T-distributed stochastic neighbor embedding (t-SNE) plots were generated based on selected PCA dimensions and unsupervised graph-based clustering was used to partition cells into clusters based on their transcriptomes. Each dot corresponds to one individual cell. Marker genes were identified by Seurat function FindAllMarkers.

#### Melanoma datasets

A single cell RNA-seq dataset of CD45+ cells from melanoma tumors was downloaded from gene expression omnibus (GEO; GSE120575) ^7^. Data were loaded into a Seurat object and clustered using the associated Seurat packages. Cells in T cell clusters with >1 CD8A transcript, >1 CD3E transcript and <2 CD4 transcripts were isolated in silico for further analysis. These cells were again clustered, and dimensionality reduction was performed using the Seurat packages. Frequency of CD8 T cells in the two clusters with the highest IL2 signaling scores (see below) were compared in patients who responded or did not respond to therapy. Wilcoxon ranked sum test was used to assign statistical significance between response groups.

#### HPV-associated cancer datasets

Single cell RNA-seq dataset of HPV-tetramer sorted CD8 T cells from HPV associated malignancies was downloaded from GEO (GSE180268) ^16^. Doublets were removed using the scds package. Low quality cells that contained >7% mitochondrial genes, <700 detected genes or <2,500 unique molecular identifiers were removed. Data from different patients were integrated using the linear regression and the batchelor package. Further clustering and dimensionality reduction was performed using the Seurat package.

#### Lung cancer patient samples

Gene expression matrices and TCR VDJ sequences from T cells isolated from non-small cell lung cancer tumors were downloaded from GEO (GSE176021) ^21^. Gene expression data for each patient were loaded into Seurat objects (version 4.1) and integrated using IntegrateData and associated functions in Seurat. Antigen specificity was assigned by mapping VDJ sequences of known neoantigen or viral specificity to gene expression data with matching cell barcodes for each patient. Low quality cells (>7% mitochondrial genes or <700 features) were removed. CD4, CD8A and CD8B gene expression was imputed using the SAVER package (version 1.1) and used to select for CD8 T cells. Subsequent clustering and dimensionality reduction were performed using the Seurat package.

#### Gene set scoring

Gene set signatures were scored using the UCell package (version 1.99). The IL2 signature gene set (FUNG_IL2_SIGNALING_1) and STAT5 targets downstream of IL-2 gene set (FUNG_IL2_TARGETS_WITH_STAT5_BINDING_SITES_T1) were downloaded from the molecular signatures database. The exhaustion and memory scores were compiled from overlapping genes in previously published datasets. Wilcoxon signed-rank tests were used to assign statistical significance.

#### Code availability

Code used to analyze human single cell RNA-seq data included R version 4.1.2 and the following publicly available packages: AUCell version 1.16 for ranking gene expression, SAVER version 1.1 for CD4, CD8A and CD8B gene imputation, scds version 1.10 for doublet detection, Seurat version 4.1, scater version 1.22 and scran version 1.22 for data manipulation and plotting, Monocle version 2.22 for pseudotime analyses and UCell version 1.99 for gene set scoring. Full code used to analyze human scRNA-seq datasets is available at https://github.com/thpulliam/IL2-exhaustion

### Statistics

Paired or unpaired Student’s t-test was used as indicated to evaluate differences between sample means. Paired analysis was conducted in experimental settings where WT and CD25KD CD8 T cells were cotransferred into the same host. Unpaired t-test was used to compare sample means when WT and KD or CD25^Hi^ and CD25^Lo^ cells were transferred into separate hosts, or to compare sample means of treated and untreated groups from *in vitro* or *in vivo* studies. A one-way analysis of variance (ANOVA) with Tukey’s post-hoc test was used for experiments with 3 or more groups to compare means across distinct groups. All statistical analysis was conducted using GraphPad Prism 5 Software (GraphPad Software, Inc.). P values of statistical significance are depicted by asterisk per the Michelin guide scale: * (P ≤ 0.05), ** (P ≤ 0.01), and *** (P ≤ 0.001) in the figures. In cases where statistically significant differences were not observed, no asterisk marks are included.

## ACKNOWLEDGMENTS

The authors would like to thank Shruti Bhise, Heather Maylor-Hagen and Yevgeniy Yuzefpolskiy for technical assistance.

## Data Availability

All data associated with this study are in the paper or supplementary materials. Microarray datasets are publicly available from previous studies at the National Center for Biotechnology Information GEO database under accession numbers GSE119943, GSE176021, GSE180268 and GSE120575.

