## Extended Data for "Tempered IL-2 Signals Program PD-1 Checkpoint Blockade-Responsive Stem-like Exhausted T Cells During Priming"

### Supplemental Data

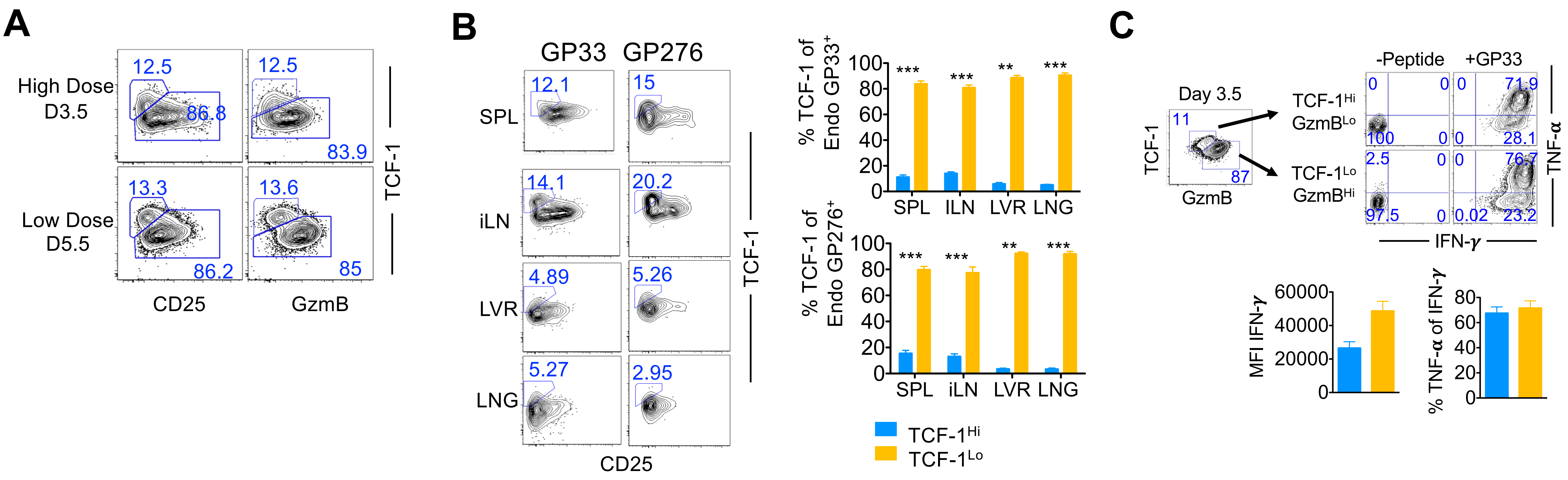


**Extended Data Fig 1. Inverse association of TCF-1 and CD25 expression is evident in TCR-transgenic CD8 T cells at low or high precursor frequencies, and in endogenous virus-specific CD8 T cells.**

**(A)** WT $D^b$GP33-specific P14 CD8 T Cells were adoptively transferred at high (1x$10^6$) or low (2.5 x$10^3$) precursor frequencies into naïve B6 mice, which were subsequently infected with $LCMV_{Cl\text{-}13}$ for 3.5 or 5.5 days respectively. Representative flow cytometry plots of TCF-1 and CD25 or TCF-1 and GzmB expression in P14 CD8 T cells on the indicated days post $LCMV_{Cl13}$ infection are presented.

**(B)** Representative flow cytometry plots show TCF-1 and CD25 expression on endogenous Gp33- and Gp276-specific CD8 T cells in the SPL, iLN, LVR and LNG at day 5.5 after infection. Bar graphs depict relative proportions of stem-like TCF-$1^{Hi}$ CD8 T cells (Light blue) or terminally differentiated TCF-$1^{Lo}$ CD8 T cells (Gold) in endogenous LCMV $D^b$GP33-, and $D^b$GP276-specific CD8 T cells in spleen at day 5.5 after infection.

**(C)** WT P14 cells (1x$10^6$) were adoptively transferred into naïve B6 recipient mice and then infected with $LCMV_{Cl\text{-}13}$ 1 day later. Representative flow cytometry plots of TNF-α+ and IFN-γ+ co-expression on gated TCF-$1^{Hi}$ $GzmB^{Lo}$ or TCF-$1^{Lo}$ $GzmB^{Hi}$ donor CD8 T cells at day3.5 post-infection are presented. Bar graphs depict MFI of IFN-γ and %TNF-α+ of IFN-γ+ in TCF-$1^{Hi}$ (Light blue) or TCF-$1^{Lo}$ (Gold) P14 T cells. Bar graphs display mean and SEM. Paired or unpaired student t-test was used with statistical significance in difference of means represented as ** ($P \leq 0.01$), *** ($P \leq 0.001$).

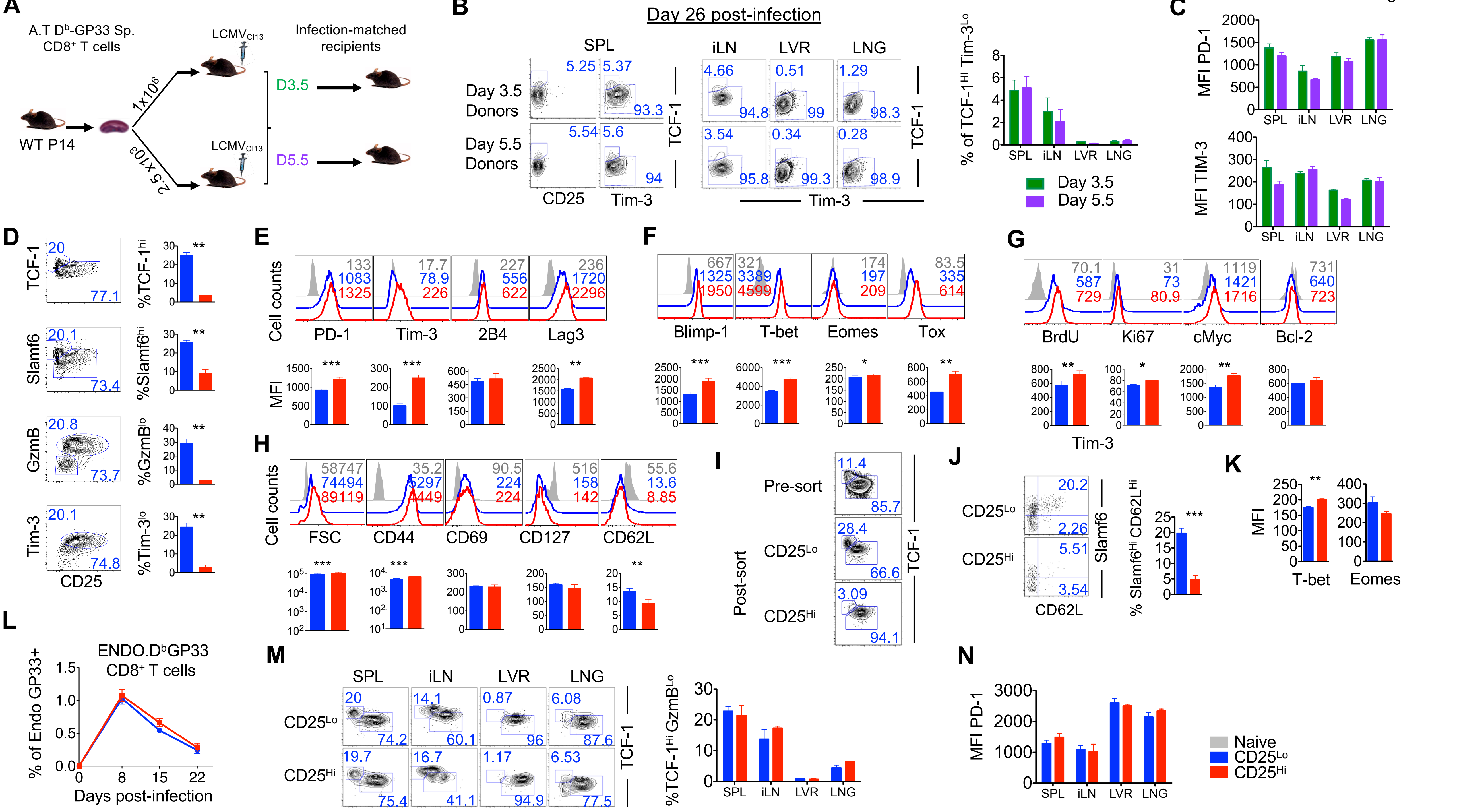
A
A.T D^b-GP33 Sp. CD8+ T cells
WT P14
1x10^6
2.5x10^3
LCMVCl13
LCMVCl13
Infection-matched recipients
D3.5
D5.5
B
Day 26 post-infection
SPL
iLN
LVR
LNG
Day 3.5 Donors
Day 5.5 Donors
CD25
Tim-3
TCF-1
% of TCF-1Hi Tim-3Lo
Day 3.5
Day 5.5
C
MFI PD-1
MFI TIM-3
D
TCF-1
Slamf6
GzmB
Tim-3
CD25
%TCF-1hi
%Slamf6hi
%GzmBlo
%Tim-3lo
E
Cell counts
PD-1
Tim-3
2B4
Lag3
MFI
F
Blimp-1
T-bet
Eomes
Tox
G
BrdU
Ki67
cMyc
Bcl-2
H
FSC
CD44
CD69
CD127
CD62L
I
Pre-sort
Post-sort
CD25Lo
CD25Hi
J
Slamf6
CD62L
% Slamf6Hi CD62LHi
K
T-bet
Eomes
L
ENDO.D^bGP33 CD8+ T cells
% of Endo GP33+
Days post-infection
M
%TCF-1Hi GzmBLo
N
MFI PD-1
Naive
CD25Lo
CD25Hi

**Extended Data Fig 2. Supplemental data related to CD25 FACS purification and adoptive transfer studies.**
**(A-C) Similar developmental profiles of adoptively transferred virus-specific CD8 T cells purified from low or high precursor frequencies.**
**(A)** Experimental setup. Chimeric C57Bl/6 mice were adoptively transferred with high dose ($1x10^6$) or low dose ($2.5 x10^3$) WT P14 CD8 T cells, and infected with $LCMV_{Cl13}$ for 3.5 or 5.5 days respectively. On the indicated days, donor cells were purified, and equal numbers of antigen specific CD8 T cells ($4x10^5$) were adoptively transferred into infection-matched recipients.
**(B)** Representative flow cytometry plots of TCF-1 with CD25 or Tim-3 in the SPL or TCF-1 and Tim-3 expression in donor CD8 T cells at day 26 post $LCMV_{Cl13}$ infection in the indicated tissues are presented. Bar graphs show the % of TCF-$1^{Hi}$ Tim-$3^{Lo}$ CD8 T cells in the indicated tissues at day 26 post-infection).
**(C)** Quantification of MFI of PD-1 and Tim-3 expression on donor CD8 T cells in the indicated tissues at day 26 post-infection, purified and transferred at day 3.5 (Green) or day 5.5 (Purple) from high dose or low dose P14 chimeric mice, respectively, are presented as bar graphs. Paired and unpaired student t-test was used to compare statistical significance in difference of means.
**(D-H) Phenotypic properties of of $CD25^{Hi}$ and $CD25^{Lo}$ virus-specific CD8 T cells.**
**(D)** Flow-cytometry plots show CD25 and TCF-1, Slamf6, GzmB or Tim-3 co-expression in splenic P14 CD8 T cells at day 5.5 post-infection. Bar graphs depict the frequency of TCF-1Hi, Slamf6Hi, GzmBLo and Tim-3Lo P14 T cells on gated CD25Lo (Blue) or CD25Hi (Red) P14 T cells 5.5 days after infection. Data presented are representative of 2 independent experiments with at least 3 mice per group.
**(E-H)** Paired histograms and bar graphs show expression of the indicated markers on gated CD25Lo (Blue) or CD25Hi (Red) antigen specific CD8 T cells in spleen; grey histograms show marker expression on endogenous CD44Lo naïve CD8 T cells. Numbers represent MFI of expression of respective markers. Data are representative of 4 independent experiments (mean ± SEM) with at least 3 mice per group.
**(I-K) Characterization of FACS sorted $CD25^{Hi}$ and $CD25^{Lo}$ at day 3.5 post infection.**
**(I)** Flow cytometry plots show TCF-1 and Tim-3 or TCF-1 and Slamf6 co-expression in splenic P14 CD8 T cells at day 3.5 post-infection before and after FACS sort.
**(J)** Representative flow cytometry plots of Slamf6 and CD62L expression on $CD25^{Hi}$ and $CD25^{Lo}$ CD8 T cells in the spleen at day 24 post infection are presented. Bar graphs depict % $Slamf6^+$ $CD62L^+$ CD8 T cells from all mice.
**(K)** Bar graphs depict the expression of T-bet and Eomes in FACS-purified and transferred $CD25^{Hi}$ (Red) or $CD25^{Lo}$ (Blue) P14 CD8 T cells at day 8 post-infection. Bar graphs display mean and SEM.
Paired and unpaired student t-test was used with statistical significance in difference of means represented as* ($P \leq 0.05$), ** ($P \leq 0.01$), *** ($P \leq 0.001$).
**(L-M) The adoptively transferred $CD25^{Lo}$ and $CD25^{Hi}$ donors minimally impact the endogenous virus-specific CD8 T cell responses.**
**(L)** Quantification of MFI of PD-1 and Tim-3 expression on donor CD8 T cells in the indicated tissues at day 26 post-infection, purified and transferred at day 3.5 (Green) or day 5.5 (Purple) from high dose or low dose P14 chimeric mice, respectively, are presented as bar graphs. Paired and unpaired student t-test was used to compare statistical significance in difference of means. **(L)** Longitudinal analysis of endogenous $D^b$GP33-CD8T cell expansion and contraction in blood at indicated times after infection.
**(M)** Representative flow cytometry plots of TCF-1 and GzmB expression in endogenous $D^b$GP33-CD8T cell-specific cells in the indicated tissues 24 days after infection. Bar graphs depict % endogenous stem-like TCF-$1^{Hi}$ $GzmB^{Lo}$ cells in $D^b$GP33-specific CD8 T cells.
**(N)** Quantification of PD-1 expression on endogenous $D^b$GP33-CD8T cells in the indicated tissues at day 24 post-infection is summarized as bar graphs. Data representative of at least 4 mice per group (mean ± SEM). Paired and unpaired student t-test was used to compare statistical significance of difference in means as appropriate. Statistical significance in difference of means is represented as * ($P \leq 0.05$), ** ($P \leq 0.01$), *** ($P \leq 0.001$).

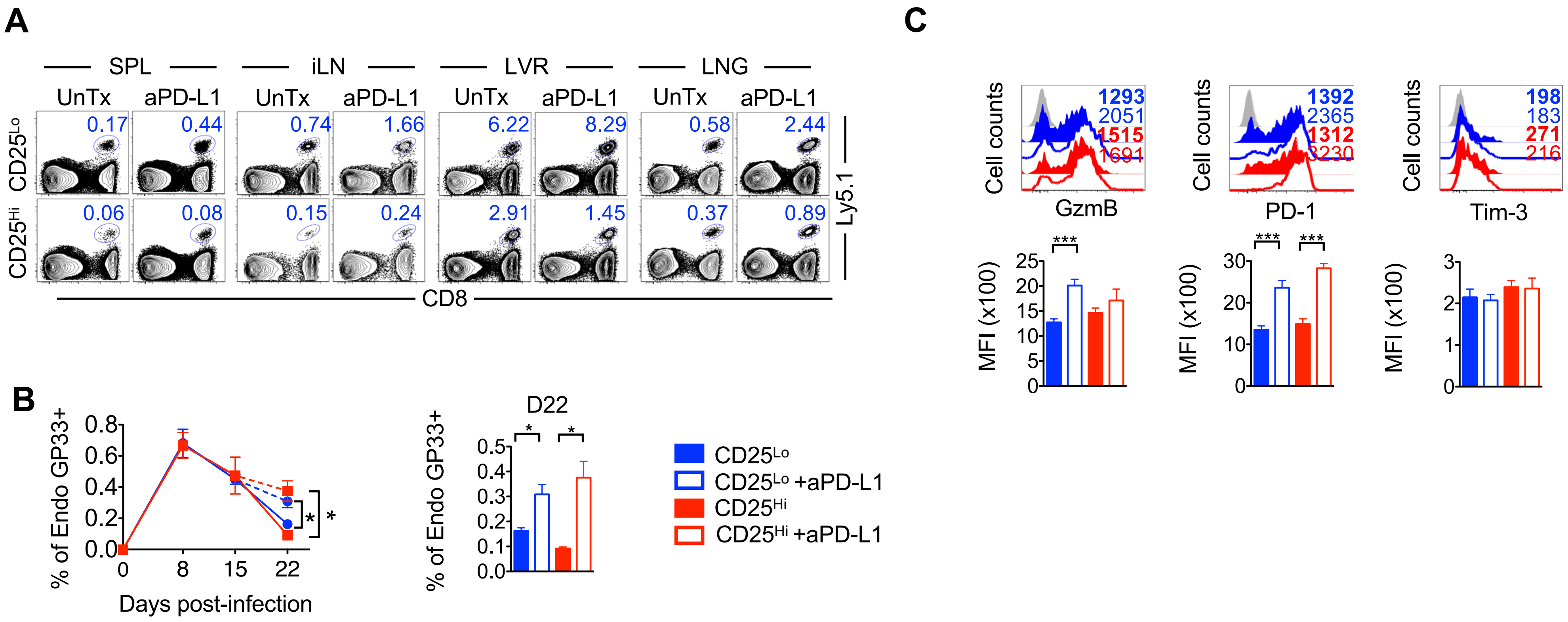


**Extended Data Fig 3. Endogenous virus-specific CD8 T cell responses to PD-1 checkpoint blockade immunotherapy remain unaltered in chronically infected recipients of CD25$^{Lo}$ and CD25$^{Hi}$ donor cells.**

**(A)** CD25Lo and CD25Hi cells were FACS purified at day 3.5 after infection, and adoptively transferred into infection-matched congenically distinct recipient mice (as in Fig 3). Mice were subsequently treated with anti-PD-L1 antibody (200 µg) every 3 days from day 15 to 24. Representative flow-cytometry plots of donor cells at day 24 after infection in the indicated tissues.

**(B)** Longitudinal analysis of endogenous DbGP33-CD8T cell expansion and contraction in blood at indicated times after infection are presented. Bar graph depicts endogenous DbGP33-specific CD8 T cell frequencies at day 22 post-infection.

**(C)** Histograms and bar graphs show GzmB, PD-1 and Tim-3 expression in endogenous D$^{b}$GP33-specific CD8 T cells in the spleen at day 24 post infection. Number represents MFI of respective marker. Bar graphs display mean and SEM. Paired and unpaired student t-test was used to compare difference of means between two groups. To compare differences between multiple groups one-way ANOVA with Tukey post-test was used with statistical significance in difference of means represented as * (P ≤ 0.05), ** (P ≤ 0.01), *** (P ≤ 0.001).

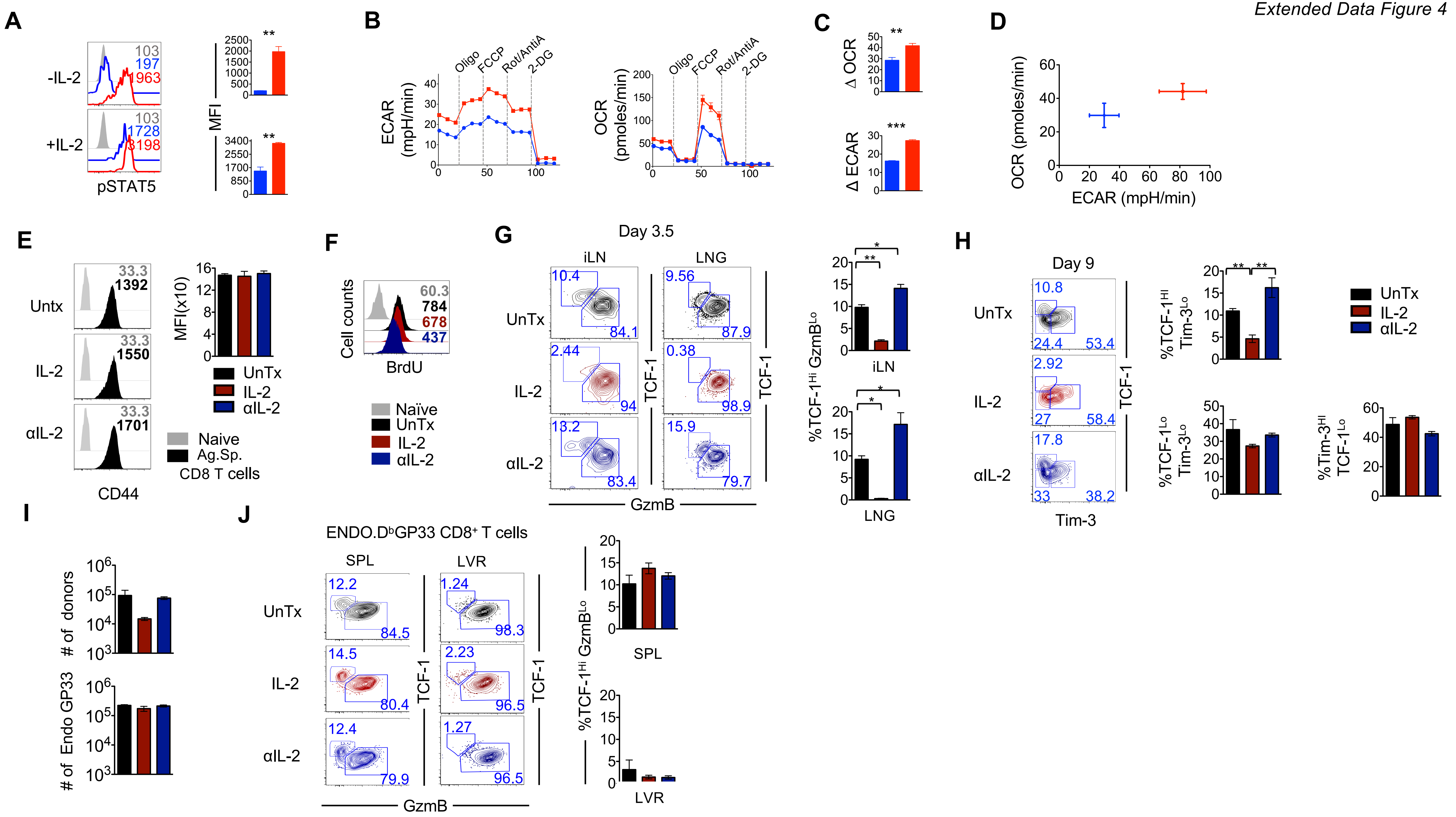
A
-IL-2
+IL-2
103
197
963
103
1728
3198
pSTAT5
MFI
B
ECAR (mpH/min)
OCR (pmoles/min)
Oligo
FCCP
Rot/AntiA
2-DG
C
Δ OCR
Δ ECAR
D
OCR (pmoles/min)
ECAR (mpH/min)
E
Untx
IL-2
αIL-2
33.3
1392
33.3
1550
33.3
1701
CD44
MFI(x10)
UnTx
IL-2
αIL-2
Naive
Ag.Sp.
CD8 T cells
F
Cell counts
60.3
784
678
437
BrdU
Naïve
UnTx
IL-2
αIL-2
G
Day 3.5
iLN
LNG
UnTx
IL-2
αIL-2
10.4
84.1
9.56
87.9
2.44
94
0.38
98.9
13.2
83.4
15.9
79.7
TCF-1
GzmB
%TCF-1Hi GzmBLo
iLN
LNG
H
Day 9
UnTx
IL-2
αIL-2
10.8
24.4
53.4
2.92
27
58.4
17.8
33
38.2
TCF-1
Tim-3
%TCF-1Hi Tim-3Lo
%TCF-1Lo Tim-3Lo
%Tim-3Hi TCF-1Lo
UnTx
IL-2
αIL-2
I
### of donors
### of Endo GP33
J
ENDO.DbGP33 CD8+ T cells
SPL
LVR
UnTx
IL-2
αIL-2
12.2
84.5
1.24
98.3
14.5
80.4
2.23
96.5
12.4
79.9
1.27
96.5
TCF-1
GzmB
%TCF-1Hi GzmBLo
SPL
LVR

**Extended Data Fig 4. Tempered IL-2 signals at priming lead to enhanced development of TCF1$^{Hi}$ CD8 T cells.**
**(A-D) Differential IL-2 signals and metabolic programming in CD25$^{Lo}$ and CD25$^{Hi}$ virus-specific CD8 T cells at priming**
**(A)** WT P14 cells ($2.5x10^3$) were adoptively transferred into naïve B6 mice, which were subsequently infected with LCMV$_{Cl13}$. Splenocytes were isolated from P14 chimeric mice at day 5.5 after LCMV$_{Cl13}$ infection. Cells were stimulated with 10nM IL-2 and intracellular p-STAT5 was assessed by flow cytometry. Histograms are gated on CD25$^{Hi}$ (red) or CD25$^{Lo}$ (Blue) CD8 T cells.
**(B)** Metabolic analysis was performed using CD25$^{Hi}$ and CD25$^{Lo}$ FACS-purified P14 CD8 T cells at day 3.5 post-infection in the presence of 10 mM glucose with indicated additions of oliomycin (oligo), trifluoromethoxy carbonylcyanide phenylhydrazone (FCCP), rotenone and antimycin A (Rot/AntiA), and 2-deoxyglucose (2DG). Line graphs show extracellular acidification (ECAR) and oxygen consumption (OCR) rates over time. Dashed vertical lines indicate pharmacologic interventions with Oligomycin, FCCP, Rotenone/AntimycinA and 2-deoxyglucose.
**(C)** Bar graphs show maximum ECAR and basal OCR on FACS sorted CD25$^{Hi}$ or CD25$^{Lo}$ at day 3.5 post-infection.
**(D)** Energy map for purified CD25$^{Lo}$ (Blue) or CD25$^{Hi}$ (Red) D$^b$GP33-specific CD8 T cells (n=8 for each mean depicted).
**(E-H) Tempered IL-2 signals at priming lead to enhanced development of TCF1$^{Hi}$ CD8 T cells.**
**(E)** Representative histograms show CD44 expression on antigen-specific CD8 T cells (Black) on day 3.5 post-infection. Grey histograms depict endogenous CD44$^{Lo}$ naïve CD8 T cells. Numbers within histograms depict MFI of marker expression.
**(F)** Representative histograms showing BrdU incorporation by the donor cells are presented. Numbers in histograms show MFI of BrdU staining for corresponding donor plots.
**(G)** Representative flow cytometry plots of TCF-1 and GzmB expression in gated D$^b$GP33-specific P14 CD8 T cells isolated from iLN and LNG at day 3.5 post-infection are presented. Bar graphs show % stem-like TCF-1$^{Hi}$ GzmB$^{Lo}$ P14 CD8 T cells. Data are representative of 3 independent experiments with n=3 mice per group.
**(H)** Representative flow cytometry plots show TCF-1 and Tim-3 expression in donor cells isolated at day 9 after LCMV$_{Cl-13}$ infection, from recipient mice adoptively transferred with day 3.5 donor cells isolated from P14 chimeric mice that were untreated (Black), or treated with IL-2 (Red) or IL-2 blocking antibody (Blue) from days 0-3.5 post-infection. Bar graphs show the frequency of TCF-1$^{Hi}$ Tim-3$^{Lo}$, TCF-1$^{Lo}$ Tim-3$^{Lo}$, Tim-3$^{Hi}$ TCF-1$^{Lo}$ on donor CD8 T at day 9 post-infection.
To compare differences between groups one-way ANOVA with Tukey post-test was used. Statistical significance in difference of means is represented as * ($P \leq 0.05$), ** ($P \leq 0.01$), *** ($P \leq 0.001$).
**(I-J) Adoptive transfer of donor cells primed in distinct IL-2 signaling conditions does not alter the differentiation program of endogenous CD8 T cells to chronic LCMV infection.**
**(I)** Bar graphs show numbers of D$^b$GP33-specific P14 donors and endogenous CD8 T cells in spleen at day 9 after infection in recipient mice adoptively transferred with day 3.5 donor cells isolated from P14 chimeric mice that were untreated (Black), or treated with IL-2 (Red) or IL-2 blocking antibody (Blue) from days 0-3.5 post-infection.
**(J)** Representative flow cytometry plots of TCF-1 and GzmB co-expression in endogenous LCMV D$^b$GP33 -specific CD8 T cells in spleen at day 9 post-infection are presented. Bar graphs depict % stem-like TCF-1$^{Hi}$ GzmB$^{Lo}$ CD8 T cells. Data are representative of at least 3 mice per group (mean ± SEM). To compare differences between groups one-way ANOVA with Tukey post-test was used. Statistical significance in difference of means is represented as * ($P \leq 0.05$), ** ($P \leq 0.01$), *** ($P \leq 0.001$).

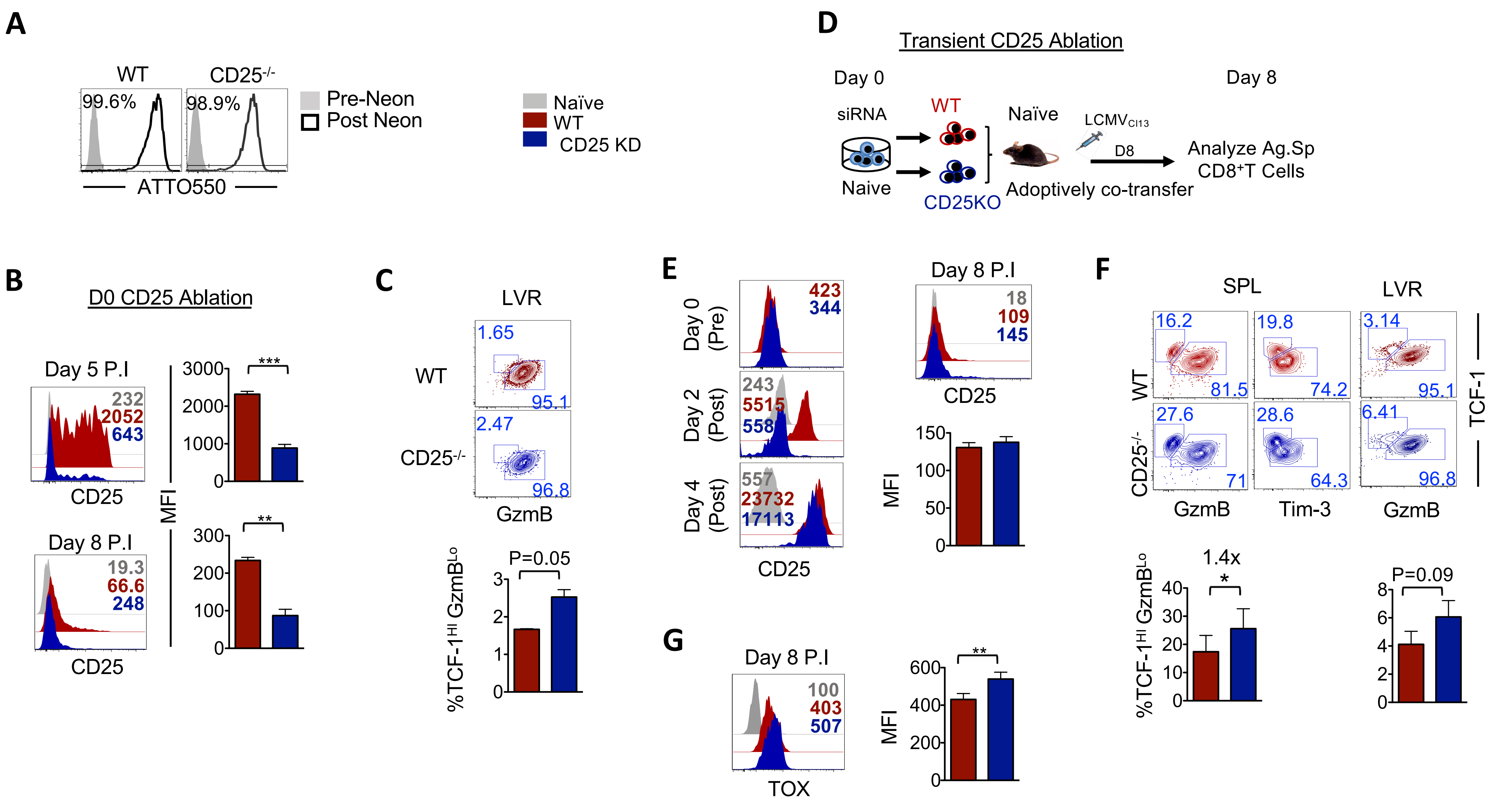
A
WT
CD25-/-
99.6%
98.9%
Pre-Neon
Post Neon
ATTO550
Naïve
WT
CD25 KD
D
Transient CD25 Ablation
Day 0
Day 8
siRNA
WT
Naïve
LCMVCl13
D8
Analyze Ag.Sp CD8+T Cells
Naive
CD25KO
Adoptively co-transfer
B
D0 CD25 Ablation
Day 5 P.I
232
2052
643
CD25
MFI
Day 8 P.I
19.3
66.6
248
CD25
C
LVR
WT
1.65
95.1
CD25-/-
2.47
96.8
GzmB
%TCF-1HI GzmBLo
P=0.05
E
Day 0 (Pre)
423
344
Day 2 (Post)
243
5515
558
Day 4 (Post)
557
23732
17113
CD25
Day 8 P.I
18
109
145
CD25
MFI
F
SPL
LVR
WT
16.2
81.5
19.8
74.2
3.14
95.1
CD25-/-
27.6
71
28.6
64.3
6.41
96.8
GzmB
Tim-3
GzmB
TCF-1
%TCF-1HI GzmBLo
1.4x
P=0.09
G
Day 8 P.I
100
403
507
TOX
MFI

**Extended Data Fig 5. Crispr/Cas9 mediated knockdown of CD25 expression in antigen-specific CD8 T cells promotes the development of TCF-1$^{Hi}$ stem-like CD8 T cells.**
**(A)** Electroporation efficiency using the Neon Transfection System. Histogram plots show percent positive cells for ATTO$^{TM}$ 55O (Black) post electroporation of D2 activated CD8 T cells.
**(B)** Paired histograms and bar graphs show the expression of CD25 on WT P14 CD8 T cells transfected with scrambled (WT) or *il2ra* guide RNAs (CD25 knockdown, CD25 KD) prior to adoptive transfer into naive mice, which were infected LCMV$_{Cl-13}$. Data are presented for WT and CD25 KD gated donor cells from spleens at day 5 and day 8 post-infection. Numbers within histograms represent MFI.
**(C)** Representative flow cytometry plots of TCF-1 and GzmB expression in WT and CD25 KD donor CD8 T cells isolated from the LVR at day 8 post-infection. Bar graphs depict composite data for % stem-like TCF-1$^{Hi}$ GzmB$^{Lo}$ CD8 T cells in gated donor populations from n=3-5 mice per group.
**(D)** Experimental setup. Naïve P14 CD8 T cells were electroporated with short interfering RNA (siRNA) targeting CD25 mRNA (CD25 KD) or with controls (WT). Equal numbers of WT or CD25 KD donor P14 CD8 T cells were adoptively co-transferred into naïve mice, which were subsequently infected with LCMV$_{Cl-13}$. Donor antigen specific CD8$^+$ T were analyzed at day 8 post-infection.
**(E)** The efficiency of siRNA-mediated transient silencing of CD25 was confirmed by flow cytometry after 48hr of *in vitro* stimulation by plate bound anti-CD3 and anti-CD28. Cells were kept in culture for two more days to confirm the re-expression of CD25 on CD25 KD cells. CD25 expression levels were also assessed in WT and CD25 KD cells adoptively transferred into C57Bl/6 mice 8 days following infection with LCMV$_{Cl-13}$. Histograms show the expression of CD25 in WT (Red), CD25 KD (Blue) and naïve CD8 T cells (Gray). Corresponding bar graphs depict MFI of CD25 expression from n=3 mice.
**(F)** Representative flow cytometry plots of TCF-1 and GzmB or TCF-1 and Tim-3 co-expression on donor CD8 T cells isolated from spleens at day 8 after infection are presented. Bar graph depicts % stem-like TCF-1$^{Hi}$ GzmB$^{Lo}$ donor CD8 T cells in spleen and liver.
**(G)** Paired histogram and bar graphs show the expression levels of TOX on WT CD8 T cells and CD25 KD donor CD8 T cells isolated from spleens at day 8 post-infection. Data are representative of at least 2 independent repeats with n=3 mice per group (mean ± SEM). Paired Student t-test was used with statistical significance in difference of means represented as * ($P \leq 0.05$), ** ($P \leq 0.01$), *** ($P \leq 0.001$).

*Extended Data Figure 6*

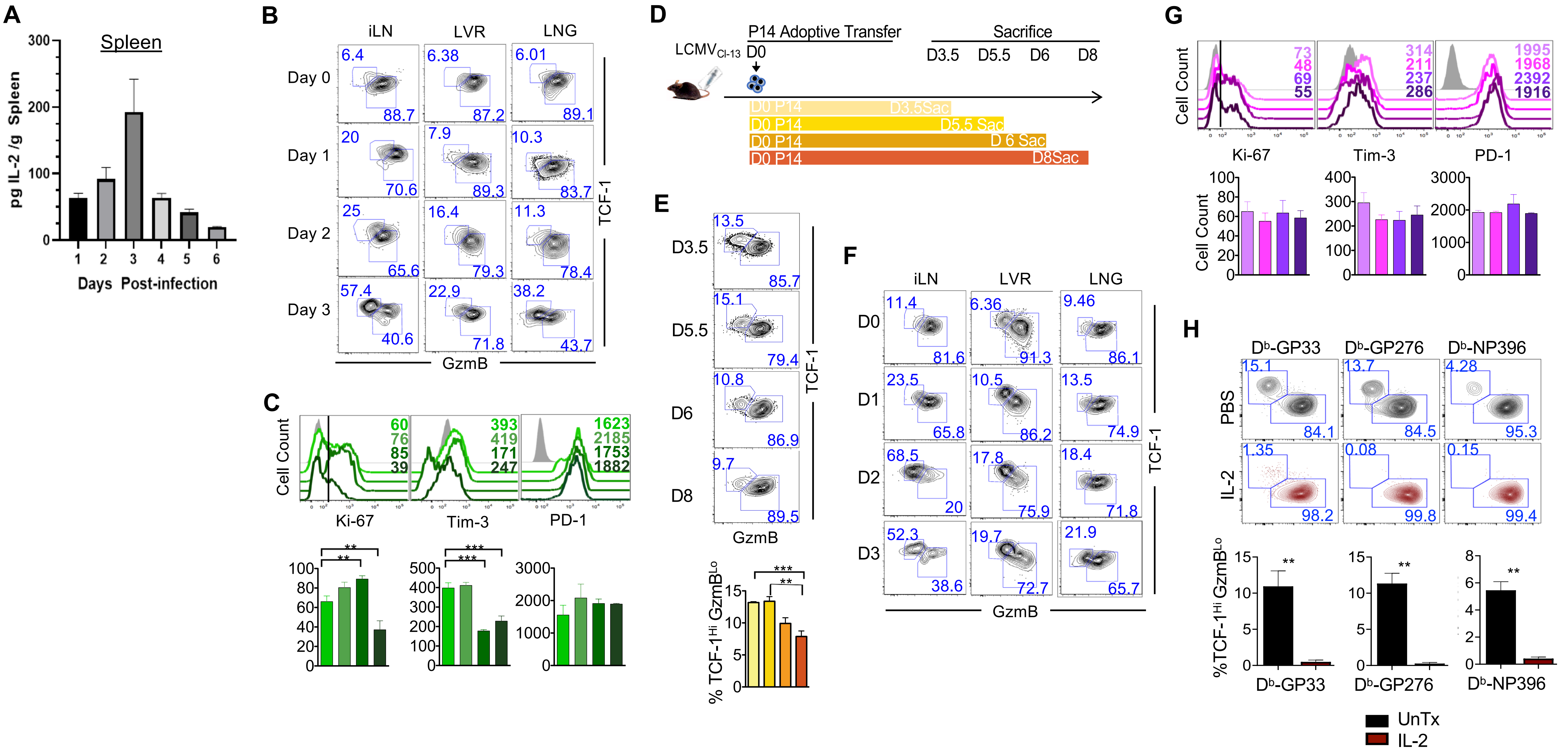

**Extended Data Fig 6. Delayed priming and curtailed duration of stimulation of virus-specific CD8 T cells promote the development of TCF-$1^{Hi}$ stem-like CD8 T cells.**
**(A)** IL-2 levels in spleen at indicated days after infection with $LCMV_{Cl-13}$ were assessed by Legendplex.
**(B)** Representative flow cytometry data for TCF-1 and GzmB expression in adoptively transferred donor CD8 T cells in iLN, LVR and LNG are presented. Virus-specific donor CD8 T cells were analyzed at day 8.5 post-infection.
**(C)** Representative histogram plots and corresponding bar charts show expression of the indicated markers on adoptively transferred $D^bGP33$-specific donor CD8 T cells in spleens at 8.5 days after stimulation; grey histograms show marker expression in endogenous $CD44^{Lo}$ naïve CD8 T cells. Numbers within histograms represent MFI of expression of respective markers. Data are representative of 2 independent experiments with n=3 mice per group.
**(D)** Experimental setup. $LCMV_{Cl13}$ infected B6 mice were adoptively transferred with $2.5x10^3$ of WT $D^bGP33$-specific P14 CD8 T cells at day 0 after infection. Donor CD8 T cells were analyzed at the indicated days post-infection in the spleens of infected mice.
**(E)** Representative flow cytometry plots of TCF-1 and GzmB expression on donor CD8 T cells in the SPL at the indicated days post-infection are presented. Bar graphs depict % TCF-$1^{Hi}$ $GzmB^{Lo}$ donor CD8 T cells in the spleen at indicated days post-infection. To compare differences between groups one-way ANOVA with Tukey post-test was used. Data are representative of 2 independent experiments with n=3 mice per group. Statistical significance in difference of means is represented as * (P ≤ 0.05), ** (P ≤ 0.01), *** (P ≤ 0.001).
**(F)** $LCMV_{Cl13}$ infected B6 mice were adoptively transferred with $2.5x10^3$ of WT P14 at days 0, 1, 2 or 3 after infection. Donor CD8 T cells were analyzed 5.5 days after P14 transfer. Representative flow cytometry plots show TCF-1 and GzmB expression on donor CD8 T cells isolated from spleens at day 5.5 post P14 adoptive transfer in the indicated tissues.
**(G)** Representative histogram plots and bar charts show levels of expression of the indicated markers on $D^bGP33$-specific CD8 T cells in spleens 5.5 days after adoptive transfer; grey histograms show marker expression in endogenous $CD44^{Lo}$ naïve CD8 T cells. Numbers within histograms represent MFI of expression of respective markers. Data are representative of 2 independent experiments with n=3 mice per group.
**(H)** P14 CD8 T cells were adoptively transferred into B6 mice at D0 or D3 after $LCMV_{Cl-13}$ infection, which were then treated with PBS or IL-2. P14 donor and endogenous GP33, GP276 and NP396 CD8 T cells were analyzed for proportions of TCF-1 and GzmB to assess the proportions of stem-like and terminally differentiated lineages. To compare differences between groups one-way ANOVA with Tukey post-test was used. Statistical significance in difference of means is represented as * (P ≤ 0.05), ** (P ≤ 0.01), *** (P ≤ 0.001).

*Extended Data Figure 7*

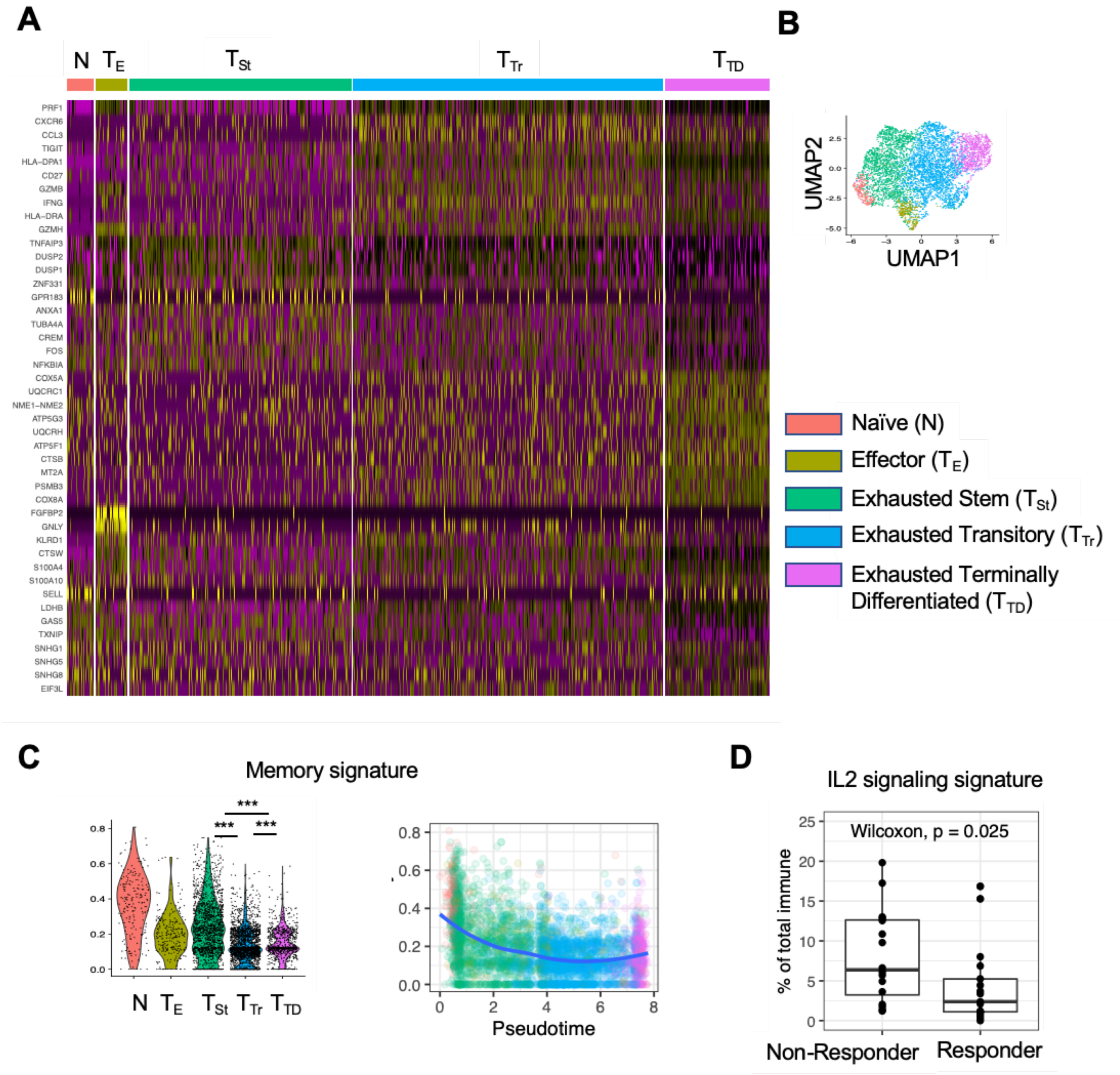

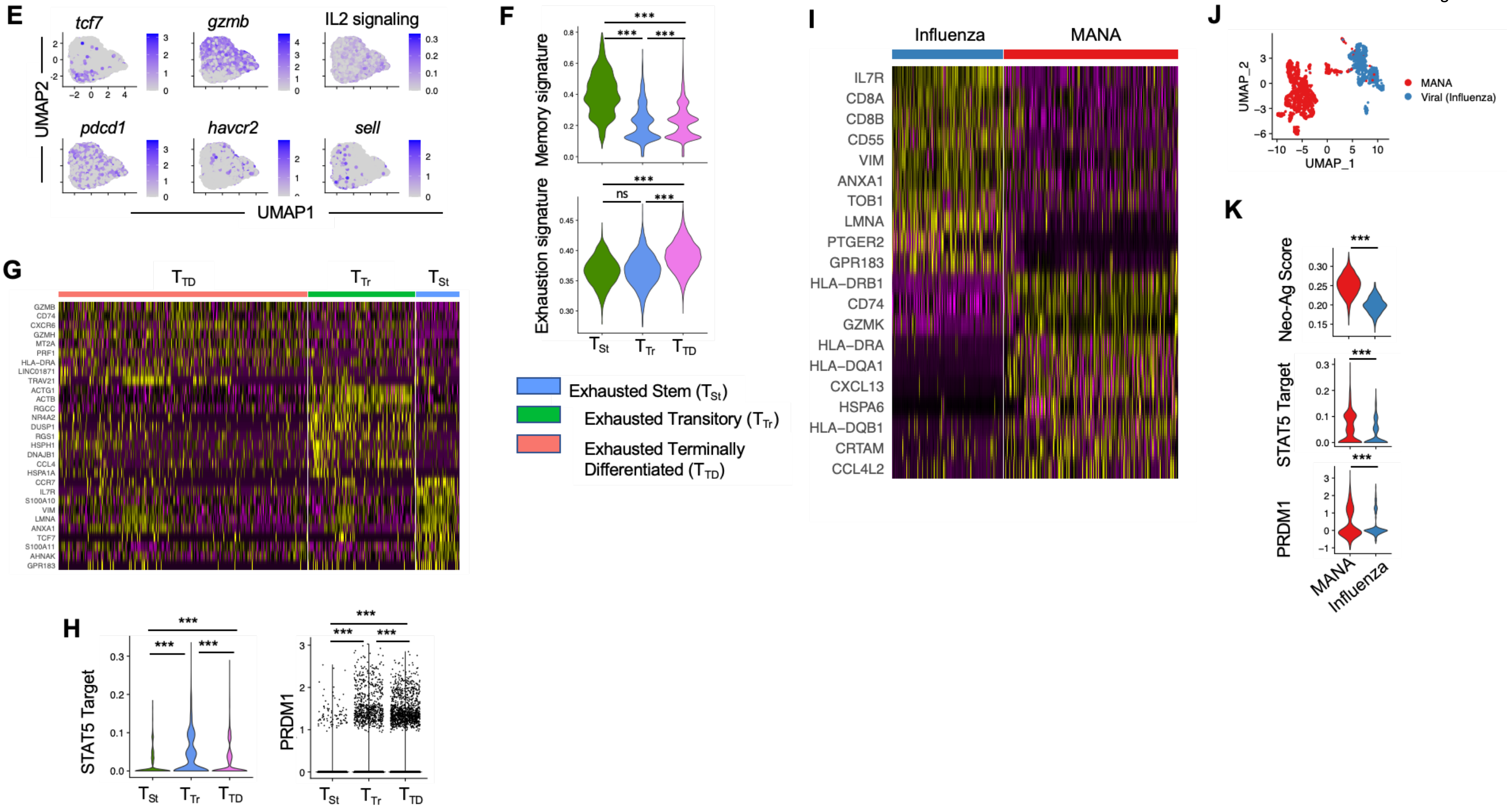
E
tcf7
gzmb
IL2 signaling
pdcd1
havcr2
sell
UMAP2
UMAP1
F
Memory signature
Exhaustion signature
ns
Exhausted Stem ($T_{St}$)
Exhausted Transitory ($T_{Tr}$)
Exhausted Terminally Differentiated ($T_{TD}$)
G
$T_{TD}$
$T_{Tr}$
$T_{St}$
GZMB
CD74
CXCR6
GZMH
MT2A
PRF1
HLA-DRA
LINC01871
TRAV21
ACTG1
ACTB
RGCC
NR4A2
DUSP1
RGS1
HSPH1
DNAJB1
CCL4
HSPA1A
CCR7
IL7R
S100A10
VIM
LMNA
ANXA1
TCF7
S100A11
AHNAK
GPR183
H
STAT5 Target
PRDM1
I
Influenza
MANA
IL7R
CD8A
CD8B
CD55
VIM
ANXA1
TOB1
LMNA
PTGER2
GPR183
HLA–DRB1
CD74
GZMK
HLA–DRA
HLA–DQA1
CXCL13
HSPA6
HLA–DQB1
CRTAM
CCL4L2
J
UMAP_2
UMAP_1
MANA
Viral (Influenza)
K
Neo-Ag Score
STAT5 Target
PRDM1
MANA
Influenza

**Extended Data Fig 7. scRNA-seq analysis of TILs from cancer patients.**
**(A-D) scRNA-seq analysis of TILs from melanoma patients.**
**(A)** Heatmap of differentially expressed genes in 5 clusters from CD8 T cells in melanoma tumors from checkpoint blockade immunotherapy treated patients. Top differentially expressed genes in each of the 5 clusters are shown on each row. Columns represent individual cells grouped by the clusters they are classified into from data set in Sade-Feldman et al., 2018.
**(B)** UMAP plot colored by clusters. Clustering shown in panel a was used to color cells in the same UMAP as in Figure 7a.
**(C)** Expression of memory signature in clusters and across pseudotime.
**(D)** Association between immunotherapy response and IL2 signature scores in CD8 T cells. The frequency of CD8 T cells from either of the clusters with high IL2-signatures (transitory $T_{Tr}$ and terminally differentiated $T_{TD}$) as a portion of all CD45+ cells were compared between patients who responded or did not respond to checkpoint blockade. Wilcoxon ranked-sum test used to establish statistical significance.
**(E-H) scRNA-seq analysis of TILs from head and neck cancer patients.**
**(E)** Expression of key exhaustion and memory genes and IL2 signaling score in HPV-specific CD8 T cells in HPV associated tumors from 15 patient dataset from Eberhardt et al., 2021. UMAP plots of scRNAseq from CD8 T cells that bound HPV-MHC tetramers CD8 T illustrating activation/exhaustion associated genes (*pdcd1, havcr2, gzmb*), memory associated genes (*tcf7, sell*) or a composite IL2 signature (FUNG_IL2_SIGNALING_1).
**(F)** Memory and exhaustion signature score in 3 clusters of HPV-specific cells from tumors. Signature scores of exhaustion and memory genes scored across Stem-like ($T_{St}$), Transitory ($T_{TR}$), and Terminally differentiated ($T_{TD}$) clusters.
**(G)** Heatmap of differentially expressed genes in 3 clusters from HPV-specific CD8 T cells. Top differentially expressed genes in each of the 2 clusters are shown in each row. Columns represent individual cells grouped by the clusters they are classified into.
Expression of IL2 signature genes and gene encoding Blimp-1 (*prdm1*) in HPV-specific CD8 T cells.
**(H)** Genes downstream of IL2 signaling that also contain STAT5 binding sites (FUNG_IL2_TARGETS_WITH_STAT5_BINDING_SITES_T1 ) were used to calculate the STAT5 Target Binding score.
**(I-K) scRNA-seq analysis of TILs from lung cancer patient tumor samples.**
**(I)** Heatmap of differentially expressed genes in T cells of various specificities in NSCLC. CD3 T cells from 15 non-small cell lung cancer (NSCLC)-tumors were isolated and single cell RNAseq was performed on these cells in Caushi et al., 2021. In parallel the MANAFEST and viraFEST assays were performed to identify mutation associated neoantigens (MANA) or influenza specific T cell receptors. CD8 T cells with either MANA or influenza-specific T cells were isolated in silico. Top differentially expressed genes for each specificity are shown in each row. Columns represent individual cells grouped by the T cell specificity based on their individual TCR sequences.
**(J)** UMAP plot of all CD8 T cells of known specificity in NSCLC. UMAP dimensionality reduction was performed on CD8 T cells of known specificity isolated in silico. Lower panel colored by exhaustion signature score.
**(K)** Violin plots of cancer-specific neoantigen STAT5 target and Blimp-1 (encoded by prdm1) scores in CD8 T cells of known specificity in NSCLC. Cancer-specific neoantigen score was derived from Lowery et al., 2022. MANA and Influenza specific CD8 T cells were isolated in silico as in panel a. Wilcoxon ranked-sum tests were used to assign statistical significance. Statistical significance represented as * ($P \leq 0.05$), ** ($P \leq 0.01$), *** ($P \leq 0.001$).
